# Dynamic fibroblast-immune interactions shape wound healing after brain injury

**DOI:** 10.1101/2024.03.13.584873

**Authors:** Nathan A. Ewing-Crystal, Nicholas M. Mroz, Anthony A. Chang, Eric Dean Merrill, Sofia E. Caryotakis, Leon Teo, Amara Larpthaveesarp, Tatsuya Tsukui, Aditya Katewa, Remy Pennington, Gabriel L. McKinsey, Sophia Nelson, Agnieszka Ciesielska, Madelene W. Dahlgren, Helena Paidassi, Saket Jain, Manish K. Aghi, James A. Bourne, Jeanne T. Paz, Fernando F. Gonzalez, Dean Sheppard, Anna V. Molofsky, Thomas D. Arnold, Ari B. Molofsky

## Abstract

Fibroblasts coordinate the response to tissue injury, directing organ regeneration versus scarring. In the central nervous system (CNS), fibroblasts are uncommon cells enriched at tissue borders, and their molecular, cellular, and functional interactions after brain injury are poorly understood. Here we define the fibroblast response to sterile brain damage across time and space. Early pro-fibrotic myofibroblasts infiltrated CNS lesions and were functionally and spatially organized by fibroblast TGF*β* signaling, pro-fibrotic macrophages and microglia, and perilesional brain glia that activated TGF*β* via integrin *α*_v_*β*_8_. Early myofibroblasts subsequently transitioned into a variety of late states, including meningeal and lymphocyte-interactive fibroblasts that persisted long term. Interruption of this dynamic fibroblast-macrophage-glial coordination impaired brain wound healing and the resolution of neuroinflammation, disrupted generation of late *de novo* CNS lymphocyte niches, and increased mortality in a stroke model. This work highlights an unexpected role of fibroblasts as coordinate regulators of CNS healing and neuroinflammation after brain injury.

---

Central nervous system (CNS) injuries including stroke, traumatic brain injury (TBI), and spinal cord injury are leading causes of death and disability,^1–3^ with chronic sequalae of cognitive/functional impairment,^4^ seizures,^5^ and neurodegeneration (e.g. Alzheimer’s Disease^6^). Treatments for CNS injuries represent a key unmet clinical need,^2,3,7^ reflecting a critical gap in our understanding of the mechanisms that dictate CNS healing.^8^ While extensive work has focused on astrocytic gliosis after injury, the CNS contains additional stromal cells, including fibroblasts, recently implicated in injury^9–13^ and disease.^14–18^ Fibroblasts are present in every organ, displaying both tissue-specific and universal/border-enriched states.^19^ Peripheral organ fibroblasts play myriad roles, mediating immune cross-regulation,^20–22^ often involving perivascular fibroblasts,^23^ or driving wound contraction, ECM deposition,^24^ and maladaptive fibrosis, often reinforced by high bioavailability of the cytokine TGFβ.^25,26^ However, the cellular origins, spatiotemporal coordination, transcriptional state(s), and functional contributions of brain fibroblasts after injury are not well understood.

## Identification of a dynamic and persistent fibroblast response to CNS injury

CNS fibroblasts are an uncommon cell type implicated in maintaining meningeal border structure, promoting immune surveillance,^27^ and regulating CSF/interstitial fluid exchange.^14^ To assess the microanatomic topography of brain fibroblasts relative to other CNS cells such as astrocytes, we first used Collagen 1 lineage tracing to specifically and indelibly mark fibroblasts (Col1a2^CreER^; R26^TdT^). In the resting brain, fibroblasts were absent from the parenchyma and localized to border regions, including larger perivascular, meningeal, and periventricular spaces, as described previously^14,28^ (**Fig. 1a, Extended Data Fig. 1a**). In a photothrombotic (PT) brain injury model involving cortical ischemia induced by a photoreactive dye, fibroblasts expanded into damaged brain regions, produced collagens and ECM, and formed a lesion distinct from parenchymal astrocytic gliosis by 14 days post injury (dpi) (**Fig. 1b**). We observed similarly widespread fibroblast infiltration in the controlled cortical impact (CCI) model of traumatic brain injury (TBI), and in a model of ischemic stroke with focal ischemia-reperfusion injury (transient middle cerebral artery occlusion, tMCAO) (**Fig. 1c,d**). In a non-human primate cortical stroke model, we also observed peri-lesional fibrosis (COL6^+^) that emerged by 7dpi and persisted through one year post injury (**Fig. 1e**). To define the kinetics of fibroblast expansion, we used orthogonal fibroblast reporter lines (e.g., Cola1^GFP^, Pdgfr*α*^GFP^) and focused on the spatiotemporally reproducible PT injury model. Fibroblast infiltration was gradual: fibroblasts were first detected near the leptomeninges by 4dpi (**Extended Data Fig. 1b**) and had surrounded and infiltrated the lesion by 14-21dpi (**Fig. 1f, Supplementary Video 1**), where they persisted for at least one year post injury (**Fig. 1g, Extended Data Fig. 1c**). Nuclear flow cytometry, which optimized fibroblast recovery, revealed that they were ∼40% of all lesional nuclei at 14dpi, a similar frequency to resting dural meninges (**Extended Data Fig. 1d,e**). We also observed injury-induced lesional vascular remodeling (**Extended Data Fig. 1f,g, Supplementary Video 2**). These findings reveal a pattern of injury-driven brain fibroblast infiltration that is coordinated, long-lasting, and conserved across damage models.

**Fig. 1:**
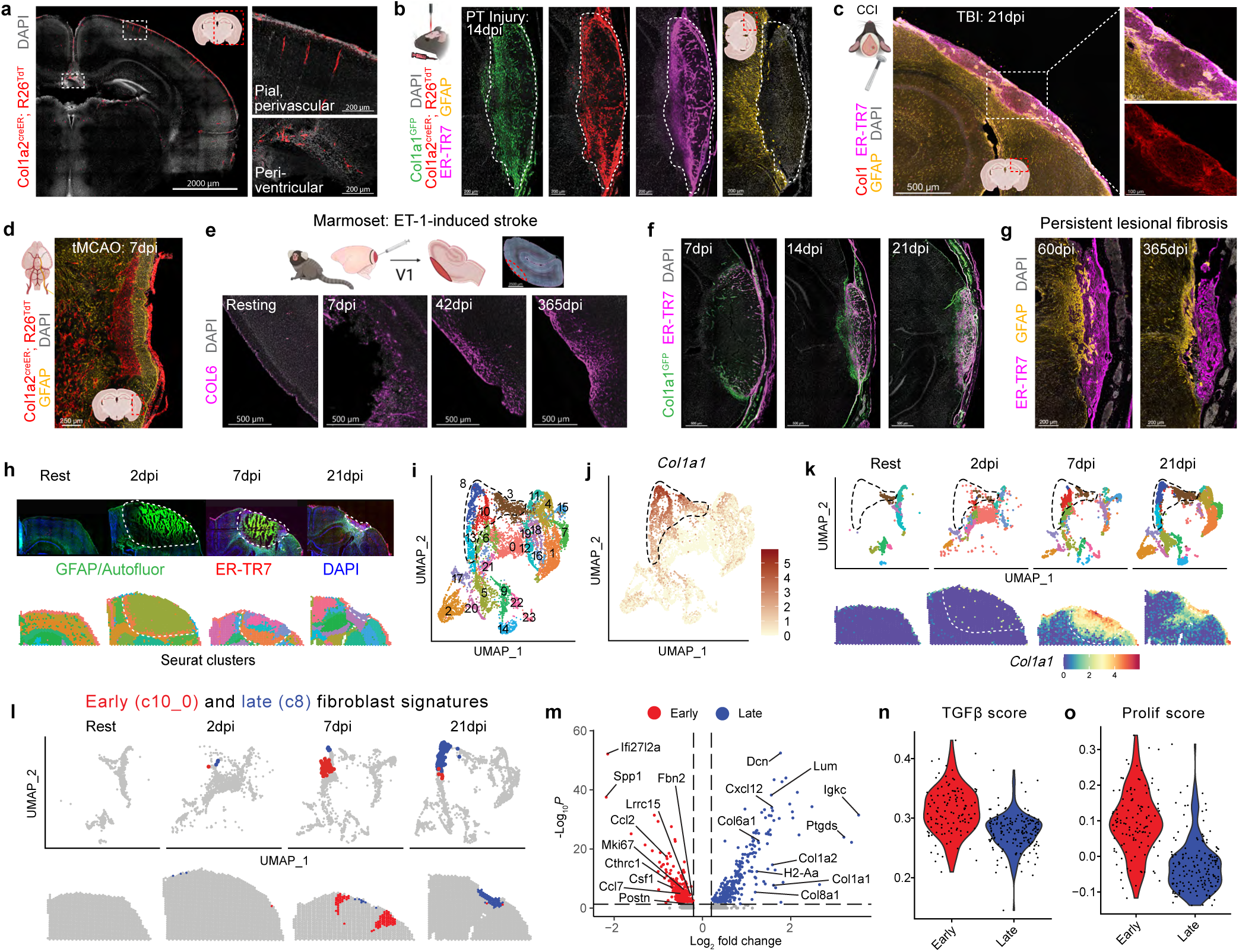
Identification of a dynamic and persistent fibroblast response to CNS injury. **a**, Homeostatic brain fibroblasts (Col1a2^creER^; Rosa26^TdT+^). Insets: pial/perivascular fibroblasts (top), periventricular/choroid plexus fibroblasts (bottom). Tamoxifen days ^-^9-^-^7 before sacrifice. **b**, Lesional fibroblasts (Col1a1^GFP+^; Col1a2^creER^; Rosa26^TdT+^) after photothrombotic (PT) injury (image of GFAP^+^ gliosis from separate mouse). Tamoxifen days ^-^16-^-^14 before injury. **c**, Fibrotic lesion (Col1^+^, ER-TR7^+^) within glial scar (GFAP^+^) after controlled cortical impact (CCI) model of traumatic brain injury (TBI). **d**, Lesional fibroblasts after transient Middle Cerebral Artery Occlusion (tMCAO) stroke. Tamoxifen days ^-^7-^-^5 before injury. **e**, Time course showing Collagen 6 (COL6) expression and persistence after endothelin-1-induced (ET-1) ischemic stroke (adult marmosets). **f-g**, PT injury time course showing fibroblasts (Col1a1^GFP+^) and/or associated ECM (ER-TR7). **h**, Immunofluorescence (top) showing astrocyte-(GFAP^+^) and fibroblast-dense regions (ER-TR7^+^), and spatial plots (bottom) with spot-based clusters. Early necrosis-associated autofluorescence is prominent; white dotted lines denote lesion borders (10X Genomics, Visium). **i-j**, UMAP plots showing spot-based clusters (**i**) or fibroblast-enriched spots (**j**, *Col1a1*^+^). 4 main fibroblast-enriched clusters highlighted (black dotted line). **k**, Time course showing fibroblast-enriched spots in UMAP (top) or spatial plots (bottom). **l-m,** UMAP and spatial plots (**l**) and volcano plot (**m**) comparing 7dpi “early-fibroblast-enriched” signature (cluster 10_0, red) and 21dpi “late-fibroblast-enriched” signature (cluster 8, blue). **n-o**, TGFβ score (**n**, genes upregulated in adventitial fibroblasts cultured with TGFβ) or proliferation score (**o**, 23 gene signature) across early/late fibroblast signatures. 200μm (**a**), 14μm (**b,f-g**), 30μm (**c**), 50μm (**d**), 40μm (**e**), or 10μm (**h**) slices; images represent two or more mice.

Next we sought to characterize PT lesional fibroblast ontogeny (i.e., cellular origin), inducing *Col1a2* lineage tracer mice prior to injury while also tracking active *Col1*-expressing fibroblasts (Col1a1^GFP^; Col1a2^creER^; R26^TdT^). By 21dpi, over half of lesional fibroblasts (Col1a1^GFP+^) were TdT^+^, indicating that they were derived from pre-existing *Col1a2*-lineage^+^ fibroblasts (**Extended Data Fig. 1h**). Lesional fibroblasts expressed canonical fibroblast markers (e.g., Gli1^TdT^, Dcn, Col6a1, Postn), while expression of mural cell markers (e.g., desmin) was sparse (**Extended Data Fig. S1i,j**). Fibroblasts did not derive from vascular smooth muscle, as determined via lineage tracing (Acta2^creER^; R26^TdT^, **Extended Data Fig. 1k**, and validating our tamoxifen washout period). To determine whether lesional fibroblasts derived from pre-existing <u>meningeal fibroblasts</u>, we applied 4-OHT, the bioactive metabolite of tamoxifen, to the intact cranium of *Col1a2* lineage tracer mice (**Extended Data Fig. 1l**). This induced sparse, local TdT expression in dural fibroblasts (**Extended Data Fig. 1m**) but not brain perivascular or leptomeningeal fibroblasts (**Extended Data Fig. 1n,o**). We observed an equal frequency of TdT^+^ fibroblasts in the dura and lesion after PT injury (∼10%, **Extended Data Fig. 1p,q**), consistent with a significant dural fibroblast contribution to lesional fibroblasts.

To molecularly define the brain fibroblast response to injury, we performed spatial transcriptomics across PT injury timepoints, capturing acute, subacute, and chronic phases of brain injury and repair (**Fig. 1h**). Dimensionality reduction revealed 23 spot-based groups (**Fig. 1i**, **Extended Data Fig. 2a,b**), with one cluster further subclustered for heterogeneity (c10; **Fig. 1i**, red). We identified four primary fibroblast-containing spatial clusters (*Col1a1*-enriched) with distinct temporal, molecular, and microanatomical patterns (**Fig. 1j,k, Extended Data Fig. 2c,d**). Two clusters that were both <u>injury-associated</u> and <u>fibroblast-enriched</u> were an early-fibroblast-enriched cluster (c10.0; **Fig. 1i**, red) that was transiently abundant at 7dpi in perilesional regions, and a late-fibroblast-enriched cluster (c8; **Fig. 1i**, blue) that was abundant at 21dpi in the lesion core (**Fig. 1l**). We also observed a resting brain border cluster (c3; **Fig. 1i**, brown) and a late lesion border cluster apparently comprising both lesional fibroblasts and adjacent reactive glia (c13; **Fig 1i**, cyan; **Extended Data Fig. 2e-g**). To begin to investigate fibroblast state evolution, we compared differentially expressed genes from the early-and late-fibroblast-enriched clusters (c10.0, red, and c8, blue, respectively; **Fig. 1m**). The early-fibroblast signature included genes implicated in pro-fibrotic processes^19,25^ and macrophage-related chemokines/growth factors. In contrast, the late-fibroblast signature was enriched for ECM molecules, potentially reflecting increased fibroblast density by 21dpi, as well as adaptive immune-associated genes.

Many early fibroblast-expressed genes were well-described targets of TGF*β* signaling, including *Cthrc1, Lrrc15,* and *Postn*.^19,25^ TGF*β*, a pleiotropic cytokine with intersecting roles in organ physiology and immunity, is particularly critical to driving pro-fibrotic ECM-depositing fibroblasts that often express *α*SMA, here called myofibroblasts.^29^ We developed a myofibroblast-associated TGF*β* score, representing genes elevated in primary fibroblasts treated with TGF*β in vitro*, and found an elevated score within the early-fibroblast signature (**Fig. 1n, Extended Data Fig. 2h**). The early-fibroblast-signature was also enriched for proliferation-related genes, a canonical feature of pro-fibrotic fibroblasts^29^ (**Fig. 1o, Extended Data Fig. 2i**). These data suggest a dynamic PT lesional fibroblast response, with early TGF*β*-associated myofibroblast activation followed by distinct late fibroblast state(s).

## Early myofibroblasts spatiotemporally correlate with lesional profibrotic macrophages and damage-associated microglia

To define brain injury-associated fibroblasts at a cellular resolution, we performed single nuclear RNA sequencing (snRNAseq) of the entire PT lesional and perilesional cortex across the injury time course, as well as uninjured cortex and cranial meninges, with a final library of 28,187 nuclei (**Fig. 2a, Extended Data Fig. 3a**). Unsupervised clustering identified expected brain cell types including neurons, astrocytes, oligodendrocytes, endothelial, immune, and stromal cells (**Extended Data Fig. 3b-f**). Stromal cell sub-clustering identified 7 lesion-enriched fibroblast clusters, 5 fibroblast clusters enriched in the resting cranial meninges, and 1 mural cell cluster (**Fig. 2b-c, Extended Data Fig. 3g**).

**Fig. 2:**
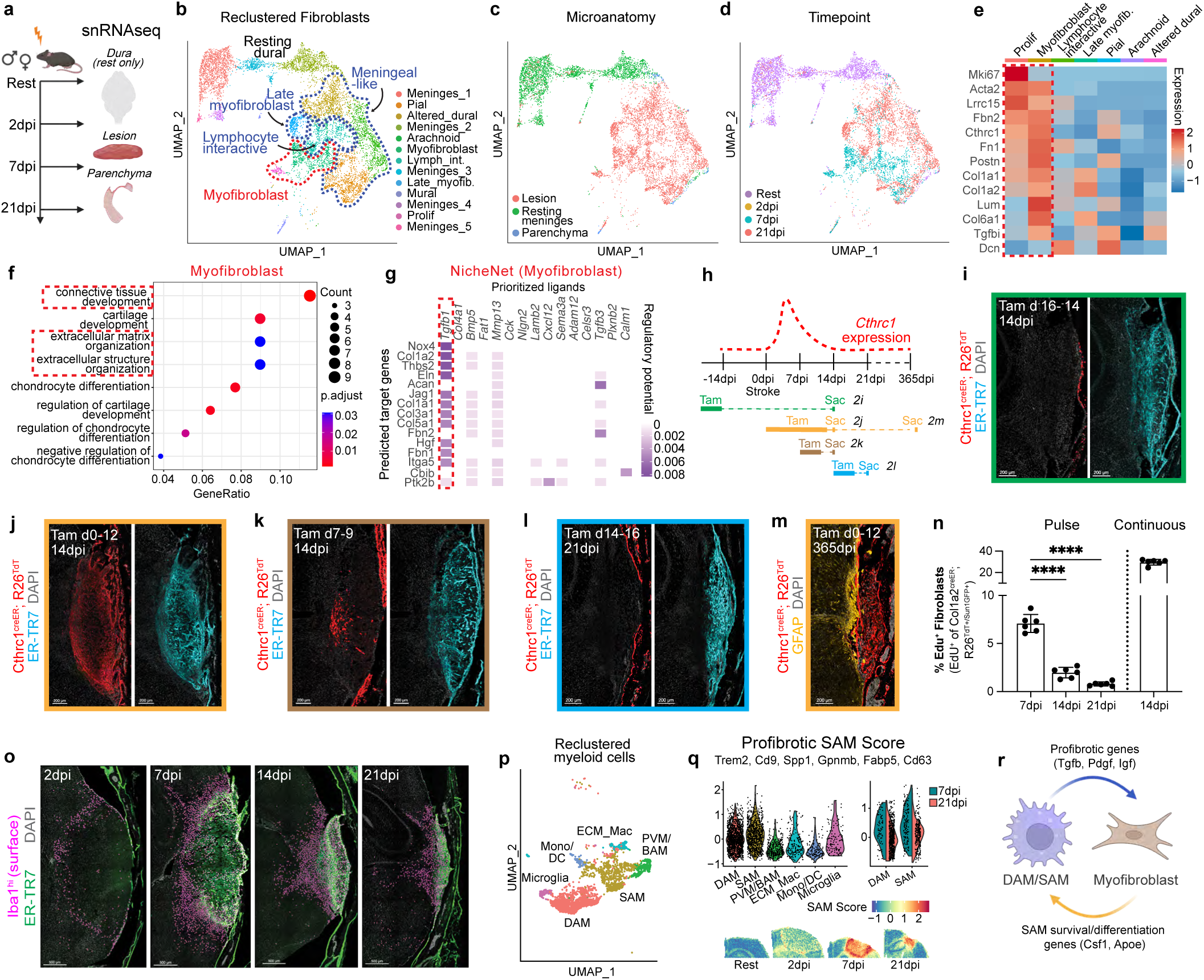
Early myofibroblasts spatiotemporally correlate with lesional profibrotic macrophages and damage-associated microglia. **a**, snRNAseq experimental schematic showing sample timepoints and microanatomy. **b-d**, UMAPs showing fibroblast subclusters (**b**), microanatomy (**d**), or timepoint (**d**). Dotted lines (**b**) highlight early myofibroblast/proliferative clusters (red) and multiple late states (blue). **e**, Heatmap of select fibrosis-related genes expressed in proliferative and myofibroblast clusters (red dotted line). **f-g**, Gene set enrichment analysis (**f**) and ligand-transcriptional-network analysis (**g**, NicheNet) within myofibroblasts. *Tgfb1* highlighted as top predicted driving ligand. **h**, Schematic showing tamoxifen induction for (**i-m**) and hypothesized *Cthrc1* expression trajectory. **i-m,** Confocal microscopy showing *Cthrc1-*lineage^+^ lesional fibroblasts after PT injury (tamoxifen induction and harvest days indicated), with lack of lineage-traced fibroblasts (**i**) but robust injury-induced *Cthrc1* (*TdT*) expression (**j**). Active *Cthrc1* expression is reduced but present at 7dpi (**k**, shown at 14dpi) and absent by 14dpi (**l**, shown at 21dpi). Labeled fibroblasts persist through 365dpi (**m**). **n**, Fibroblast proliferation (EdU incorporation) 2 hours after EdU pulse (“Pulse”; 7dpi, 14dpi, or 21dpi) or after EdU injection every other day (“Continuous”; 14dpi). Fibroblasts defined as Col1a2^creER^; Rosa26^TdT+/Sun1GFP+^. n=6 slices/timepoint (2 slices/mouse). **o**, Confocal microscopy showing time course of myeloid cell (Iba1^hi^ surface) lesional accumulation. **p-q**, Annotated UMAP of reclustered myeloid cells (**p**) and violin or spatial plots of profibrotic “SAM score” (**q**; *Trem2, Cd9, Spp1, Gpnmb, Fabp5,* and *Cd63*). In **q**: left, score by cluster; right, score by timepoint within “DAM” and “SAM” lesional clusters; bottom, score mapped onto spatial transcriptomic Visium data. **r**, Schematic showing potential ligand-receptor interactions between SAM/DAM and myofibroblasts, derived from **Extended Data Fig. 6n**. Graphs show mean±s.d. ****p<0.0001; one-way ANOVA, Tukey post-test. 14μm slices; images represent two or more mice.

Lesional fibroblasts were temporally heterogeneous, mirroring findings suggested by spatial sequencing (**Fig. 2d, Extended Data Fig. 3h**). Focusing first on early timepoints, we observed predominantly myofibroblast and proliferative myofibroblast states. Early myofibroblasts expressed canonical markers including *Cthrc1, Acta2, Lrrc15, Postn, Fn1, Tgfbi,* and *Dcn* (**Fig. 2e**), with enrichment for programs associated with ECM organization and connective tissue/cartilage development (**Fig. 2f**). Using computational prediction, *Tgfb1* was proposed as the ligand most likely to drive the early myofibroblast state (**Fig. 2g**). We observed similar populations of early injury-associated and resting fibroblasts in the non-human primate stroke model (**Extended Data Fig. 4a,b**) with clusters enriched after injury (7dpi) that expressed canonical myofibroblast genes (**Extended Data Fig. 4c-e**). Analysis of published scRNAseq data from human patients with TBI also revealed a small fibroblast cluster (**Extended Data Fig. 4f**), and post-TBI fibroblasts expressed myofibroblast-associated genes (**Extended Data Fig. 4g,h**). Using data from human glioblastoma multiforme patients, fibroblasts were also identified (**Extended Data Fig. 4i-k**); the largest fibroblast subcluster expressed myofibroblast genes (**Extended Data Fig. 4l,m**). These data are consistent with a conserved cross-species myofibroblast-like response to diverse brain injuries.

To orthogonally visualize this early myofibroblast response, we performed cellular lineage tracing for *Cthrc1*,^30^ a gene selectively expressed in pro-fibrotic fibroblasts with high levels of TGF*β*-signaling (Cthrc1^creER^; R26^TdT^, **Fig. 2h**). *Cthrc1* lineage^+^ fibroblasts were sparse in resting meninges and cortex, and these did not give rise to lesional fibroblasts (**Fig. 2i, Extended Data Fig. 5a**). However, continuous C*thrc1* labeling during PT injury revealed widespread lineage^+^ fibroblasts in the lesion and injured dural meninges (**Fig. 2j, Extended Data Fig. 5a, Supplementary Video 3**). Using timed tamoxifen induction, we found that injury-induced *Cthrc1* expression occurred between 0 and 7dpi and was fully lost by 14dpi (**Fig. 2k,l**); however, lineage-traced fibroblasts persisted for at least one year (**Fig. 2m**). Similar myofibroblast kinetics were observed using *α*SMA lineage tracer mice, which additionally labeled expected vascular smooth muscle cells (**Extended Data Fig. 5b-g**). Proliferation (EdU incorporation) was also elevated in early myofibroblasts (**Fig. 2n, Extended Data Fig. 5h**), as suggested by our transcriptomic data.

Macrophages are phagocytic immune cells associated with peripheral tissue damage and fibrotic responses.^26,31,32^ Concurrent with an early myofibroblast response, we observed an injury-responsive population of CNS myeloid cells, including reactive microglia and macrophages, that formed a perilesional ring by 2dpi, subsequently infiltrated the PT lesion core, and persisted for weeks (**Fig. 2o, Extended Data Fig. 6a**), with substantial spatial association with fibroblasts (**Extended Data Fig. 6b**). To define the ontogeny of the myeloid response, we tracked cells deriving from reactive microglia (i.e., damage-associated microglia/DAM)^33^ via P2ry12^creER^, infiltrating monocytes^34^ via Ccr2^creER^, or border associated/perivascular macrophages (BAM/PVM^32^) via PF4^cre^. Microscopy revealed that myeloid cells from both microglial and monocytic ontogenies were present at the acute lesional ring and inhabited regions of subacute emerging fibrosis (**Extended Data Fig. 6c-e**). While microglia-and monocyte-derived cells spatially diverged by 14dpi – with enrichment in the outer border and lesional core, respectively – they remained overlapping in the fibroblast-rich inner lesional border (**Extended Data Fig. 6f**). Flow cytometry confirmed that early infiltrating monocytes differentiated into lesional macrophages and dendritic cells (**Extended Data Fig. 6g,h**). In contrast, BAM/PVM-lineage macrophages were diffusely present across lesional areas and time points (**Extended Data Fig. 6i**).

We further defined injury-associated myeloid cell heterogeneity via snRNAseq, with sub-clustering identifying homeostatic microglia, damage-associated microglia (DAM), inflammatory monocytes and dendritic cells, and several macrophage subsets (**Fig. 2p, Extended Data Fig. 6j**-l). Macrophage subsets were further parsed as (a) PVM/BAM,^32^ (b) macrophages resembling scar-or lipid-associated macrophages (SAM/LAM) with pro-fibrotic roles in other organs,^26^ here named SAM, and (c) low abundance “ECM Macrophages”. A recently described profibrotic “SAM score”^26^ was highly expressed not only in SAM but also in DAM, particularly at 7dpi, and this profibrotic score, along with SAM/DAM cluster signatures, was enriched in areas of lesional fibrosis (**Fig. 2q, Extended Data Fig 6m**). To investigate potential mechanisms underlying this profibrotic state, we performed ligand-receptor analysis, which predicted that SAM/DAM myeloid cells could signal to myofibroblasts via profibrotic signals including *Tgfb1*, *Pdgfa/b/c,* and *Igf1* **(Fig. 2r, Extended Data Fig. 6n)**, and that lesional myofibroblasts could reciprocally influence SAM/DAM via *Csf1* (M-CSF) and *ApoE*, a ligand for TREM2 that promotes DAM states.^33^ Collectively, this bioinformatic and spatial analysis suggests an early and transient profibrotic identity convergence in resident microglia and recruited monocyte-derived macrophages; this convergent state is spatiotemporally associated with lesion-border myofibroblasts and may reflect a response to shared environmental cues such as dead cells and debris, lipids, and cytokines.^26^

## Distinct late fibroblast states include lymphocyte interactive fibroblasts associated with T cell accumulation and long-term persistence

While myofibroblasts predominated during early subacute phases of CNS injury, they were replaced by a collection of distinct late fibroblast states by chronic stages (21dpi, **Fig. 3a**). One cluster, lymphocyte interactive fibroblasts, expressed particularly high levels of signals for lymphocyte recruitment and support (e.g., *Cxcl12, Ccl19, Cxcl16,* and *Ccl8*),^27,35^ in contrast to early myofibroblast expression of myeloid-recruiting chemokines (e.g., *Ccl2/Ccl7*) (**Fig. 3b**). Indeed, lymphocyte interactive fibroblasts were enriched for pathways associated with the adaptive immune response (**Fig. 3c**). Mirroring the emergence of this fibroblast cluster, T lymphocytes were rare in the uninjured brain but accumulated by 14dpi (**Fig. 3d, Supplementary** Fig. 1a). Infiltrating lesional T cells were a mixture of CD4^+^ and CD8^+^, with a relative enrichment of CD8^+^ T cells at later timepoints (**Fig. 3e**); most CD4^+^ T cells expressed Tbet (*Tbx21*; **Fig. 3f, Extended Data Fig. 7a**), consistent with a T helper type 1 (Th1) IFN*γ*-expressing identity. T cells persisted through at least 60dpi (**Fig. 3d**), and lesioned hemispheres were enriched for a tissue resident memory (TRM) lymphocyte phenotype (CD44^+^ CD69^+^, **Extended Data Fig. 7b**).

**Fig. 3:**
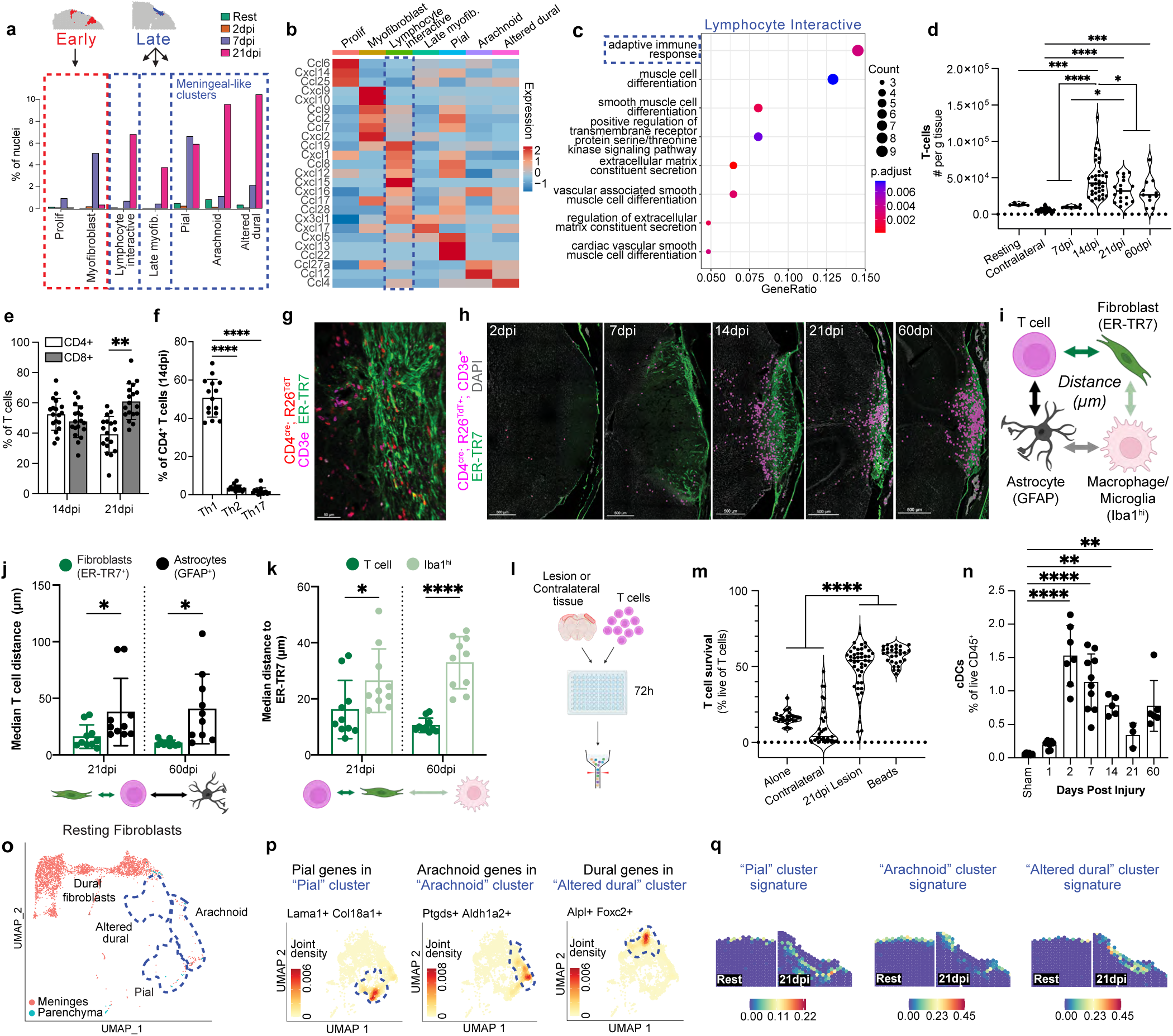
Distinct late fibroblast states include lymphocyte interactive fibroblasts associated with T cell accumulation and long-term persistence. **a**, Fibroblast cluster abundance over time, with corresponding early/late spatial signatures. **b**, Heatmap of cluster-defining chemokines, with lymphocyte interactive fibroblasts highlighted (blue dotted line). **c**, Gene set enrichment analysis among lymphocyte interactive fibroblasts. **d-f,** Total T cells (**d**, CD3^+^), CD4^+^/CD8^+^ T cells (**e**), or Th1/2/17 CD4^+^ T cells (**f**, Tbet^+^/Gata3^+^/RORγt^+^; 14dpi) after PT injury (cortical flow cytometry). n=5 (rest), 46 (contralateral), 6 (7dpi), 38 (14dpi), 18 (21dpi), or 11 (60dpi) mice (**d**); 18 mice/timepoint (**e**); or 15 mice (**f**). **g-h**, Image (**g**) and time course (**h,** surfaces) showing T cells (CD4^cre^; Rosa26^TdT+^; CD3e^+^) near fibroblast-rich lesions (ER-TR7^+^). **i**, Schematic for proximity analysis between T cells (CD4^cre^; Rosa26^TdT+^; CD3e^+^) or myeloid cells (Iba1^hi^, macrophages/reactive microglia) and fibroblast ECM (ER-TR7) or reactive astrocytes (GFAP). **j-k**, Median T cell distance from nearest fibroblast-ECM or astrocyte surface (**j**), and T or myeloid cell distance from nearest fibroblast surface (**k**). 21/60dpi. n=10 slices/timepoint (2 slices/mouse). **l-m**, Schematic for *ex vivo* lesional coculture (**l**) and quantification of T cell survival (**m**, 21dpi lesions). n=30 (alone), 31 (contralateral), 44 (21dpi lesion), or 29 (beads) wells. **n**, Conventional dendritic cell (cDC) infiltration and persistence after PT injury (cortical flow cytometry). n=6 (sham/60dpi), 7 (1/2dpi), 10 (7dpi), 5 (14dpi), or 3 (21dpi) mice. **o**, Resting fibroblasts colored by microanatomy; blue dotted lines show post-injury meningeal-like clusters. **p**, Co-expression of pial, arachnoid, and dural genes^36^ among corresponding clusters. **q**, snRNAseq signatures mapped onto spatial transcriptomic data, showing meningeal-like fibroblast distribution at rest (left, meningeal border) and 21dpi (right, new lesional-parenchymal border). Graphs show mean±s.d. *p<0.05, **p<0.01, ***p<0.001, ****p<0.0001; one-way ANOVA, Tukey post-test (**d**,**f**,**m**,**n**); multiple T-tests, Holm-Sidak correction (**e**,**j**,**k**). 14μm slices; images represent two or more mice.

To detect potential fibroblast-lymphocyte interactions, we defined spatial relationships between T cells and lesional fibroblasts, increasing our sensitivity by using T cell reporter mice (CD4^cre^; R26^TdT^, **Fig. 3g-i**). By 21dpi and through 60dpi, T cells more tightly and specifically associated with fibroblast ECM than with reactive astrocytes (**Fig. 3j, Extended Data Fig. 7d**), with fibroblast-associated T cells preferentially persisting over time (**Extended Data Fig. 7e**). T cells were closer to fibroblast-dense regions, and more lesion-associated, than total macrophages/microglia (**Fig. 3k, Extended Data Fig. 7f**), highlighting a specific spatial relationship between T cells and late lesional fibroblasts (**Extended Data Fig. 7g-l**). To directly test the ability of fibroblast-dense lesions to support lymphocyte survival, we developed an *ex vivo* assay wherein primary T cells were cocultured with dissected brain lesions (**Fig. 3l**). We found that 21dpi lesions supported increased T cell survival (**Fig. 3m, Extended Data Fig. 7m, Supplementary** Fig. 1b) without impacting T cell proliferation or activation over 72 hours (**Extended Data Fig. 7n-p**). Late 21dpi lesions promoted more T cell survival compared with early 7dpi lesions (**Extended Data Fig. 7q**). Additionally, conventional dendritic cells (cDCs), which accumulated during injury, were enriched in lesioned cortex through 60dpi (**Fig. 3n, Extended Data Fig. 7r**,s), consistent with the potential for local antigen presentation. Collectively, these data suggest the presence of late lymphocyte-interactive “niche” fibroblasts that recruit and support brain lymphocytes and their associated functions, including type 1 cytokine secretion.

A distinct fibroblast cluster enriched at 21dpi was labeled “late myofibroblasts”; these fibroblasts expressed high levels of ECM genes but lacked canonical markers of a highly active profibrotic state (e.g., *Acta2*, *Cthrc1, Lrrc15*), suggestive of diminished activation via TGF*β* or other signaling pathways (**Fig. 2e**). Instead, late myofibroblasts expressed markers associated with smooth muscle differentiation (e.g., *Cdh18, Sema3c*) and inhabited the lesional core, in contrast to border-enriched lymphocyte-interactive fibroblasts (**Extended Data Fig. 7t,u**), potentially reflecting an emergent perivascular/mural-like identity associated with vascular remodeling (**Extended Data Fig. 1f,g**). The final three late fibroblast (21dpi) clusters were identified as pial, arachnoid, and altered dural fibroblast states, based on resting leptomeningeal fibroblast distribution and meningeal layer signatures^36^ (**Fig. 3o,p**). Integration of these meningeal snRNAseq cluster identities with spatial sequencing revealed their predicted, resting border distribution, as well as the formation of a new, expanded post-injury meningeal layer between lesion and CNS parenchyma (**Fig. 3q**), recapitulating molecular and microanatomical aspects of the homeostatic meninges during wound healing.

## CNS myofibroblasts are coordinated by macrophages, perilesional glia, and TGF*β* to drive wound healing and limit chronic inflammation and mortality

We next investigated the functional impacts of CNS fibroblast-immune dialogue during injury responses. Given observed spatiotemporal associations between lesional profibrotic macrophages and myofibroblasts, we first functionally tested macrophage contributions (**Fig. 4a**). We used clodronate liposomes to deplete CNS infiltrating monocytes, early macrophages, and total perilesional myeloid cells (**Extended Data Fig. 8a-e, Supplementary Fig. 1c,d**). Clodronate liposomes reduced lesional fibroblastic ECM coverage (**Fig. 4b**) and led to increased lesion size (**Fig. 4c**), implicating macrophages in myofibroblast expansion and CNS wound healing. Ligand-receptor analysis highlighted *Tgfb1* as one macrophage ligand that could contribute to profibrotic roles (**Fig. 2r, Extended Data Fig. 6n**). Indeed, *Tgfb1* colocalized with perilesional myeloid cells by 2dpi, and myeloid cells had the highest *Tgfb1* transcript levels after injury (**Extended Data Fig. 8f,g**). We investigated the functional importance of myeloid *Tgfb1* by inducing PT injury in mice with microglia and macrophages that were induced to be TGF*β*1-deficient (Cx3cr1^creER^; Tgfb1^flox/GFP-KO^; **Extended Data Fig. 8h,i**). Conditional knockouts had lower lesional fibroblastic ECM coverage (**Extended Data Fig. 8i,j**), consistent with a partial contribution of myeloid *Tgfb1* to the injury-induced fibroblast response.

**Fig. 4:**
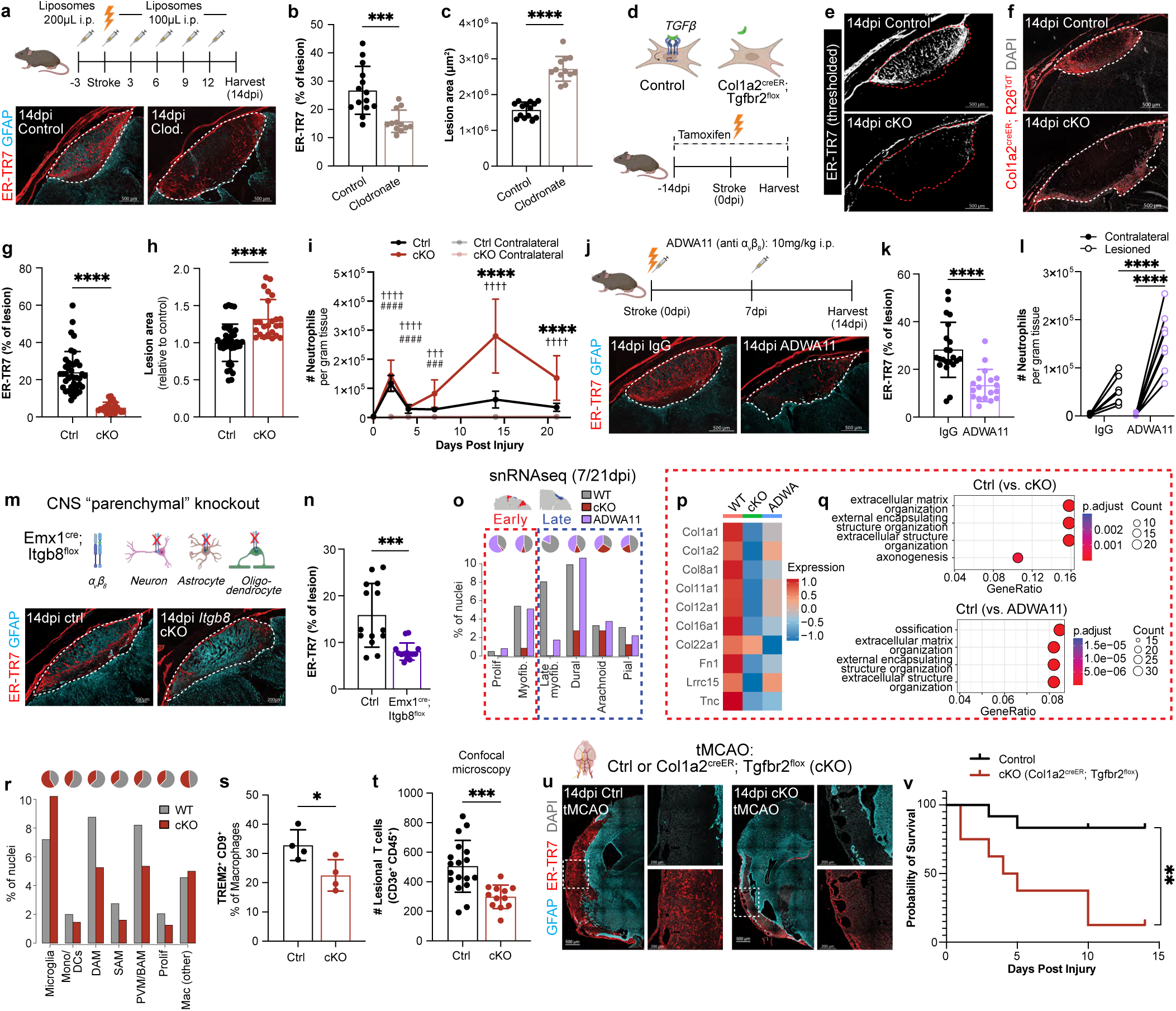
CNS myofibroblasts are coordinated by macrophages, perilesional glia, and TGFβ to drive wound healing and limit chronic inflammation and mortality. **a-c,** Lesional images after control/clodronate liposomes (**a**), ECM (ER-TR7) coverage (**b**), and lesion size (**c**), 14dpi. n=14 (control) or 12 (clodronate) slices (2 slices/mouse). **d**, Schematic showing ablated TGF*β*-signaling in Col1a2^creER^; Tgfbr2^flox^ cKO fibroblasts (tamoxifen days ^-^14-^-^12, ^-^7, 0-2, 5, 8, etc.). **e-h**, Control/cKO lesional images with thresholded ER-TR7 (**e**) or lineage traced fibroblasts (**f**, Col1a2^creER^; Rosa26^TdT+^), ER-TR7 coverage (**g**), and lesion size (**h**, normalized per experiment), 14dpi. n=40 (control) or 24 (cKO) slices (2 slices/mouse). **i**, Cortical neutrophil time course in controls/cKOs. n=6 (0dpi-control [unrelated], 2/4/7dpi-control, 2/7/14/21dpi-cKO), 8 (14dpi-control), 9 (21dpi-control), 3 (0dpi-cKO), or 5 (4dpi-cKO) mice. **j-l**, Control (IgG) or *α*_v_*β*_8_-blocked (ADWA11) lesional images (**j**), ER-TR7 coverage (**k**), and cortical neutrophils (**l**), 14dpi. n=22 (IgG) or 18 (ADWA11) slices (**k**, 2 slices/mouse); n=7 (IgG) or 8 (ADWA11) mice (**l**). **m-n**, Control or Emx1^cre^; Itgb8^flox^ lesional images (**m**) and ER-TR7 coverage (**n**), 14dpi. n=14 slices/group (2 slices/mouse). **o**, Fibroblast cluster abundance (snRNAseq) in control, cKO (Col1a2^creER^; Tgfbr2^flox^), and ADWA11-treated mice (7/21dpi). Red/blue boxes highlight early/late-enriched states. **p-q**, Myofibroblast-related heatmap (**p**) and gene set enrichment (**q**) among myofibroblasts; controls show ECM/ECM-process enrichment. **r-t**, Myeloid cluster abundance (**r**), SAM frequency (**s**, flow cytometry), or perilesional T cells (**t**) in controls/cKOs. n=4 mice/group (**s**); n=18 (control) or 12 (cKO) slices (**t**, 2 slices/mouse). **u-v**, 14dpi fibrotic lesion (**u**, ER-TR7^+^, with cKO tissue loss) or survival (**v**) after tMCAO. n=12 (control) or 8 (cKO) mice (tamoxifen days ^-^3-^-^1, 2, 5, 8, 11). Graphs show mean±s.d. *p<0.05, **p<0.01, ***p<0.001, ****p<0.0001; * lesioned-control vs. lesioned-cKO; # lesioned-control vs. contralateral-control; ^†^lesioned-cKO vs. contralateral-cKO (**i**). Student’s T-test (**b**,**c**,**g**,**h**,**k**,**n**,**s**,**t**); two-way ANOVA, Sidak’s post-test (**i** [per timepoint], **l**); Gehan-Breslow-Wilcoxon test (**v**). 14μm slices; dotted lines show lesion boundary (**a,e,f,j,m**).

We next tested the functional role of CNS myofibroblasts. TGF*β* promotes myofibroblast differentiation, activation, proliferation, migration, and ECM production in peripheral wound healing, as well as many fibrotic and inflammatory pathologies.^29^ Given the diverse roles of TGF*β* in development,^37^ we used a genetic model where fibroblasts can be “blinded” to all TGF*β* signaling in an inducible manner (Col1a2^creER^; Tgfbr2^flox^, **Fig. 4d**). In these conditional knockout (cKO) mice, brain lesional fibroblasts and associated ECM were greatly reduced (**Fig. 4e-g, Extended Data Fig. 9a**) with increased lesion size (**Fig. 4h**), suggesting impaired myofibroblast infiltration and downstream wound closure or contraction. Fibroblast topography was unperturbed in cKO uninjured tissue (**Extended Data Fig. 9b**), consistent with a less active contribution from the myofibroblast state in the resting adult brain. Flow cytometry of PT-injured cortex revealed that infiltrating neutrophils were initially elevated but declined by 4dpi in control mice; however, cKO mice had lesional neutrophilia through 21dpi (**Fig. 4i, Extended Data Fig. 9c**), reflecting persistent or secondary innate inflammation. Microscopy confirmed increased late perilesional neutrophils (**Extended Data Fig. 9d**). Lesional monocytes were also increased at late time points (**Extended Data Fig. 9e,f**). Persistent myeloid inflammation was isolated to the lesioned cortical hemisphere, as no difference in neutrophils or monocytes was found in the contralateral hemisphere, meninges, blood, or spinal cord (**Extended Data Fig. 9g-l**), nor did we detect overt spinal cord abnormalities (**Extended Data Fig. 9m**). Fibroblasts regulate the adjacent blood brain barrier,^14^ and leakage/bleeding resulting from loss of this support^12^ could contribute to neuroinflammation and tissue loss. However, while we observed increased vascular leakage/potential bleeding after injury, as determined by Evans Blue extravasation, there was no further increase in cKO mice (**Extended Data Fig. 9n**). Together, these data suggest that TGF*β*-driven myofibroblast expansion after CNS injury is critical for wound healing and the resolution of lesional innate inflammation.

TGF*β* is canonically secreted as a latent cytokine that requires activation prior to signaling.^37^ Integrins bind and activate the latent TGF*β* complex, including integrins *α*_v_*β*_1_, *α*_v_*β*_6_, and *α*_v_*β*_8_.^38^ While CNS co-expression of *α*_v_*β*_1_ or *α*_v_*β*_6_ was restricted to dural meningeal fibroblasts, *α*_v_*β*_8_ co-expression was high in astrocytes and oligodendrocytes, and variably present among lesional fibroblasts (**Extended Data Fig. 9o**). To test the potential functional role of the candidate integrin *α*_v_*β*_8_, we utilized a well-validated blocking antibody, ADWA11.^39^ Mice treated with ADWA11 phenocopied *Tgfbr2* cKO mice with reduced lesional ECM coverage (**Fig. 4j,k**), although lesion sizes were not significantly increased (**Extended Data Fig. 9p**). As in cKOs, ADWA11 drove persistent neutrophilic inflammation (**Fig. 4l**) and trending increases in monocytes (**Extended Data Fig. 9q**) that were specific to the lesioned cortex (**Extended Data Fig. 9r,s**). *Itgb8* reporter mice showed high TdT expression in perilesional GFAP^+^ astrocytes and minimal expression in lesional fibroblasts (**Extended Data Fig. 9t**). To functionally confirm the possible role of glial *α*_v_*β*_8_, we genetically deleted *Itgb8* from the brain cortical parenchyma, including glial cells (**Fig. 4m**; Emx1^cre^; Itgb8^flox^ mice). After PT injury, there was reduced lesional ECM (**Fig. 4m,n**) with unchanged lesion size (**Extended Data Fig. 9u**), phenocopying ADWA11. These data implicate lesion-adjacent brain glial cells in licensing optimal TGF*β*-mediated lesional myofibroblast expansion via *Itgb8*, although other integrins and/or mechanisms of TGF*β* activation may also contribute.^37^

We next investigated the cellular and molecular consequences of interrupting CNS fibroblast wound-healing programs. We again performed snRNAseq of PT lesional tissue and associated residual parenchyma at subacute and chronic timepoints, testing mice with TGF*β*-blind fibroblasts (cKO, Col1a2^creER^; Tgfbr2^flox^), disrupted *Itgb8*-driven TGF*β* activation (ADWA11), and controls, with a final library of 60,070 nuclei (**Extended Data Fig. 10a**). After identification of expected CNS cell types (**Extended Data Fig. 10b-d**), we focused on lesional fibroblast subsets, which mapped onto our previously defined CNS fibroblast states (**Extended Data Fig. 10e-j)**. cKO mice had a profound loss of early myofibroblast clusters at 7dpi and a substantial reduction in late states at 21dpi, including late myofibroblasts, meningeal-like, and lymphocyte-interactive fibroblasts, supporting our model of sequential state transitions (**Fig. 4o, Extended Data Fig. 10k**). ADWA11-treated mice had relatively preserved numbers of early myofibroblasts and late meningeal and lymphocyte-interactive fibroblasts, but 21dpi late myofibroblasts were nearly absent. Additionally, early myofibroblasts from both cKO and ADWA11-treated mice expressed fewer ECM and pro-fibrotic genes (**Fig. 4p,q**), consistent with a shared impairment of early myofibroblast differentiation.

Given predicted bidirectional fibroblast-macrophage interactions, we also defined impacts of cKO myofibroblast perturbation on lesional myeloid cell heterogeneity (**Extended Data Fig. 10l-n**). cKO mice had reduced DAM and SAM clusters (**Fig. 4r**), and trajectory analysis revealed impaired monocyte-to-SAM and microglia-to-DAM transitions (**Extended Data Fig. 10o,p**). We orthogonally validated a relative decrease in SAM via flow cytometry (**Fig. 4s**), supporting an interruption of reciprocal myofibroblast-profibrotic macrophage interactions (**Extended Data Fig. 10q**). ADWA11-treated mice showed an expected cross-macrophage pro-inflammatory and “dysmature” transcriptional signature, consistent with known homeostatic roles for *Itgb8*-mediated TGF*β* autocrine signaling in microglia^40,41^ (**Extended Data Fig. 10r,s**). Potentially reflecting a loss of myofibroblast-derived lymphocyte-interactive fibroblasts, we found reduced lesion-associated T cells in myofibroblast-impaired cKO mice, with residual T cells largely confined to the disrupted lesion-parenchymal interface (**Fig. 4t, Extended Data Fig. 10t**). Lymphocytes from cKO mice also showed transcriptomic differences by 21dpi, particularly within CD8^+^ and *γδ* T cell clusters (**Extended Data Fig. S10u,v**) where cKO lymphocytes expressed increased *Itgae* (CD103, **Extended Data Fig. 10w**). CD103 is a residency marker for CD8 T cells, including in the brain parenchyma,^42^ potentially reflecting a distinct perilesional lymphocyte topography in the absence of a lesional fibroblast niche. These data suggest that evolving stromal-immune interactions shape both the innate and the adaptive immune landscapes after PT brain injury.

Finally, we sought to characterize clinically relevant contributions of fibroblasts in CNS injury and repair. As PT damage is a relatively mild model with little functional brain impairment, we instead turned to tMCAO, a more severe ischemia-reperfusion injury model that mirrors aspects of human stroke microanatomy and pathophysiology,^43^ and tested mice with TGF*β*-blind fibroblasts (cKO, Col1a2^creER^; Tgfbr2^flox^). By 14dpi, control mice formed robust fibrotic lesions, whereas surviving cKO mice displayed impaired fibrotic, but not glial, scar formation as well as substantial cortical tissue loss (**Fig. 4u**). cKO mice also died at higher rates (∼85% versus ∼15% of controls, **Fig. 4v**). These data suggest that effective brain wound healing requires functional myofibroblasts, which are reinforced by transient pro-fibrotic macrophages/microglia, and more broadly highlight the potential clinical relevance of CNS fibroblast responses (**Fig. 5**).

**Fig. 5:**
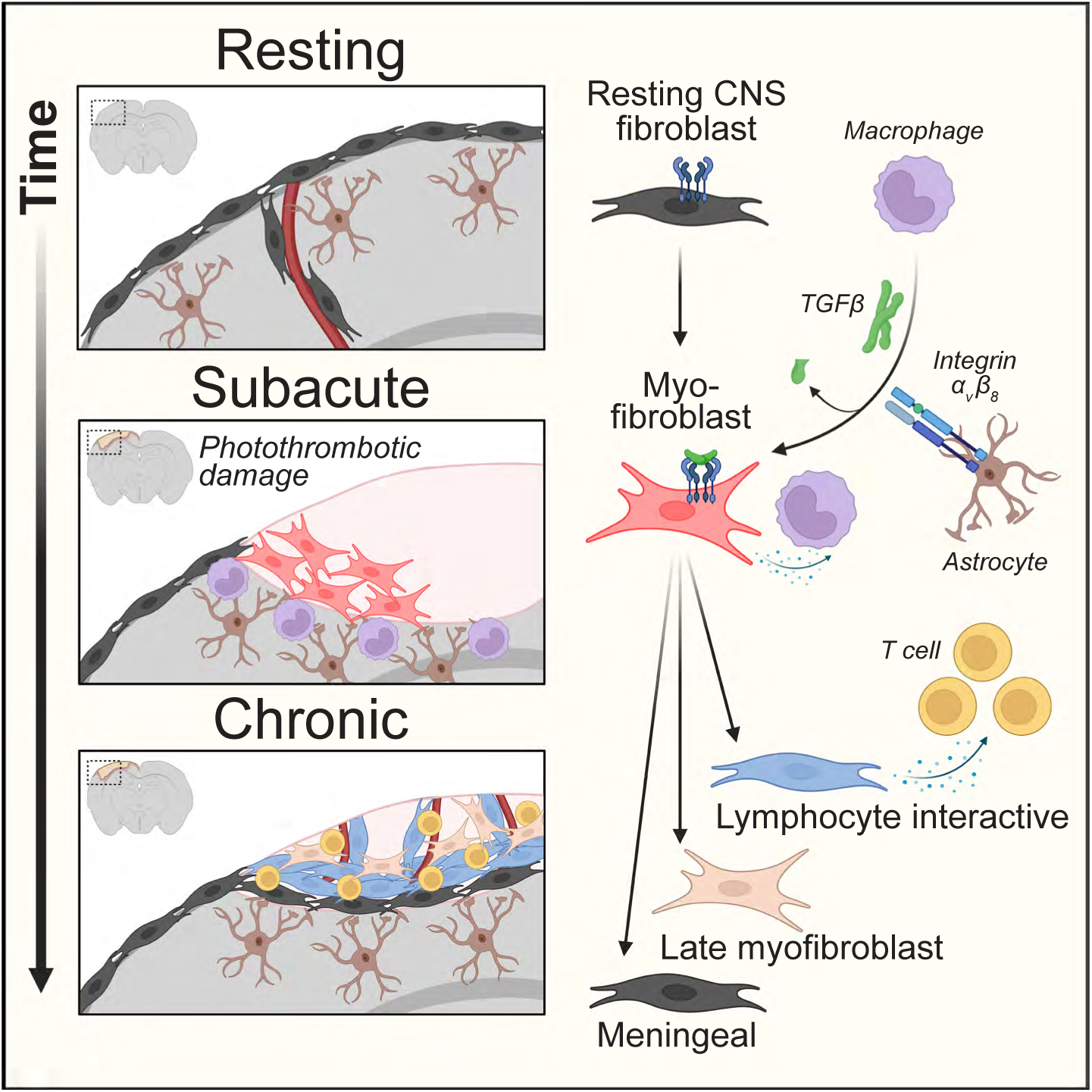
Graphical Summary.

### Discussion

Here we characterize a functionally critical fibroblast-immune conversation across sterile brain injury, conserved in several models and species, involving fibroblast activation, proliferation, migration, and ultimate persistence. While stromal cells such as fibroblasts are recently appreciated participants in CNS physiology and pathology, their ontogeny has remained unclear. Distinct studies arguing for pericyte^9–11,13^ or fibroblast^44^ origins for damage-associated CNS stromal cells have relied on nonspecific markers^14^ or qualitative lineage tracing.^12^ Additionally, in contrast to peripheral organ injury-associated fibroblasts^24,25^ or resting CNS fibroblasts,^27,36,45,46^ the states/identities of CNS fibroblasts after injury are poorly defined. We use *Col1a2* lineage tracing to quantitatively define resting CNS fibroblasts as a major source for PT injury-responsive fibroblasts. We additionally employ integrated spatial and single nuclear transcriptomics to characterize fibroblast 4D evolution. Early proliferative and profibrotic myofibroblasts, which engage in reciprocal interactions with profibrotic macrophages, subsequently transition to distinct late states including meningeal border fibroblasts and lymphocyte-interactive fibroblasts that persist long-term and appear to chronically alter the CNS immune landscape. These transitions may reflect evolving tissue requirements after injury-induced disruption of CNS anatomic and immunologic protection,^47^ with early prioritization of physical boundary reformation and later prioritization of border homeostasis, including meningeal recapitulation and lymphocyte recruitment/retention. The long-term impacts of de novo CNS lymphocyte niches also remain to be defined. Fibroblasts display context-dependent state transitions in other organs and diseases, including skin wounding,^24^ lung fibrosis,^30^ ulcerative colitis,^20^ and cancers,^19,48,49^ consistent with a role as coordinators of organ homeostasis attuned to complex and dynamic tissue needs. Notably, the brain is a sterile environment with relatively few resting fibroblasts, providing a tabula rasa that allows for dissociation of the impacts of microbial signals and/or pre-existing tissue fibroblasts abundant in other barrier sites.

We also define the TGFβ pathway as a critical activator of brain fibroblasts after injury. Disruption of TGFβ cellular sources, including convergent profibrotic macrophages and microglia, TGF*β* activators, including glial cells expressing *α*_v_*β*_8_, or TGF*β* receptors attenuated brain fibroblast responses. In mice with TGFβ-blind fibroblasts, both early myofibroblasts and late fibroblast states were ablated; consequently, PT lesions were larger with unresolving innate inflammation surpassing the post-injury peak. While the mechanisms and consequences of this inflammation, which was also observed after *α*_v_*β*_8_ blockade, remain to be fully elucidated, they may involve feedback loops that increase inflammatory damage to CNS parenchyma and drive subsequent recruitment of further innate inflammatory cells. Myofibroblasts driven by TGF*β* may prevent this cycle by forming a new emergency border, involving ECM and/or locally enriched chemokines that limit neutrophil migration, between susceptible parenchyma and ischemic tissue^50^ or disrupted vasculature.^28^ Additionally, some fibroblasts may maintain immune-suppressive identities to impact this balance.^48^ Finally, impaired fibrotic lesion formation was associated with increased cortical tissue loss and higher mortality in a severe stroke model (tMCAO). Collectively, these data reveal the critical importance of CNS fibroblast-immune dialogue in recovery after injury, extending the domain of fibroblasts as key drivers of wound healing to the CNS. Given the newly appreciated presence of stromal cells across human CNS diseases, fibroblasts may play broader roles in CNS health and pathology and represent intriguing future therapeutic targets.

## METHODS

### Mice

To lineage trace resting fibroblasts and track injury-responsive fibroblasts, we crossed Col1a2^creERT2^ mice (MGI 6721050, from Bin Zhou, Institute of Biochemistry and Cell Biology, Shanghai Institutes for Biological Sciences)^51^ with Rosa26^TdT-Ai14^ mice (R26-CAG-RFP, Jackson 007914) containing a flox-stop-flox sequence upstream of a CAG-RFP-WPRE cassette in the constitutively expressed ROSA26 locus. Tamoxifen induces RFP expression in *Col1*-lineage^+^ cells. Col1a2^creERT2^ mice were also crossed to Rosa26^Sun1GFP^ mice (Jackson 030952), enabling fibroblast nuclear identification. Additional stromal reporters used include Col1a1^GFP^ mice (from David Brenner, University of California, San Diego) to mark active *Col1-*expressing cells;^52^ Pdgfra^GFP^ (PDGFRa-H2B-eGFP nuclear-localized GFP, Jackson 007669) to track fibroblasts and oligodendrocyte lineage cells;^53,54^ and Rosa26^Tdt-Ai14^ mice crossed to Gli1^creERT2^ mice (Jackson 007913) to track adventitial fibroblasts;^23^ Twist2^cre^ mice (Jackson 008712) to track fibroblasts and mural cells;^55^ Acta2^creERT2^ mice to track myofibroblasts and smooth muscle cells;^56^ and Cthrc1^creER^ mice (generously provided by Dean Sheppard) to track myofibroblasts.^30^

To track immune cell subsets, we crossed Rosa26^TdT-Ai14^ mice to CD4^cre^ mice (Jackson 022071) track CD4^+^ and CD8^+^ T cells;^57^ Ccr2^creERT2^ mice (from Burkhard Becher, University of Zurich, Switzerland) to track monocyte-derived cells; P2ry12^creERT2^ (Jackson 034727) to track microglia-derived cells; Cx3cr1^creER^ (Jackson 020940) to track macrophages; and PF4^cre^ (Jackson 008535) to track border associated macrophages. We also used T-bet (Tbx21)-zsGreen transgenic mice (generously provided by Jinfang Zhu, Lab of Immune System Biology, NIH)^58^ to track type 1 lymphocytes. Cx3cr1^creER^ mice were additionally crossed with Tgfb1^GFP^ mice (MGI 3719583)^59^ and Tgfb1^flox^ mice (Jackson 033001) to drive deletion of *Tgfb1* in macrophages (in Cx3cr1^creER^; Tgfb1^GFP/flox^ mice). Controls were littermate Cx3cr1^creER^; Tgfb1^flox/+^ mice.

To conditionally delete fibroblast *Tgfbr2*, we crossed Col1a2^creERT^^2^ mice to Tgfbr2^flox^ mice (both Tgfbr2-exon2^flox^, MGI 2384513,^60^ and Tgfbr2-exon4^flox^, Jackson 012603); Additionally, we used Itgb8^TdT^ mice (Itgb8-IRES-TdT, generously provided by Helena Paidassi)^61^ to visualize *Itgb8* expression, and Itgb8^flox^ mice (MGI 3608910)^62^ crossed to Emx1^cre^ mice (Jackson 005628) to delete *Itgb8* from neurons and glial cells.^41,63,64^ For above conditional knockout strains, controls were littermate Cre-negative or flox-heterozygous mice.

Mice were mixed gender animals backcrossed on C57BL/6 for at least 10 generations, or on a mixed genetic background (Gli1^creERT2^, Cx3cr1^creER^). If not otherwise stated, all experiments were performed with 8-16 week old male and female mice. All mice were bred and maintained in specific-pathogen-free conditions at the animal facilities of UCSF and were used in accordance with institutional guidelines and under study protocols approved by the UCSF Institutional Animal Care and Use Comittee.

### Marmoset

n=11 outbred middle-aged marmoset monkeys (*Callithrix jacchus;* >5 years; median age ∼7 years) were used in this study. No siblings were used. Animals were housed in family groups (12:12 hrs light/dark cycle, temperature 31°C, humidity 65%). Experiments were conducted according to the Australian Code of Practice for the Care and Use of Animals for Scientific Purposes and were approved by the Monash University Animal Ethics Committee. Marmosets were obtained from the National Nonhuman Primate Breeding and Research Facility (Monash University, Australia).

### Tamoxifen-induced Cre recombination

Mice were injected i.p. with 200μL of tamoxifen (Sigma-Aldrich) dissolved in corn oil at 10mg/mL. For transcranial activation, 4-OH-tamoxifen (Sigma-Aldrich), the active metabolite of tamoxifen,^65^ was dissolved in acetone at 100μg/mL. 100μL of 4-OHT was applied to a specific cranial location, determined stereotactically as described below (“Photothrombotic injury”). Tamoxifen was applied via micropipette (approximately 5μL at a time) and allowed to evaporate in between applications.

### Photothrombotic injury

For PT injury surgeries,^66–68^ mice were anesthetized via inhaled isoflurane, shaved on the scalp, and stereotaxically fixed. After sterilization with iodine and 70% ethanol, 0.5% lidocaine was administered subcutaneously to the scalp. The cranium was surgically exposed, and a fiber optic white light is placed over the S1 cortex (3.0mm,-0.5mm (x,y) from bregma, with coordinates determined via stereotax). Mice were injected intraperitoneally with 8mg/kg Rose Bengal dye; after 1 minute, the cranium was exposed to high intensity white light (150W) for 2 minutes. The scalp was sutured using nylon sutures and surgical glue, and buprenorphine was administered.

### TBI

For TBI surgeries,^5^ mice were anesthetized and stereotaxically fixed, as above. A 3-mm craniotomy was performed over the right S1 centered at –1 mm posterior from bregma, +3 mm lateral from the midline. TBI was performed with a CCI device (Impact One Stereotaxic Impactor for CCI, Leica Microsystems) equipped with a metal piston using the following parameters: 3 mm tip diameter, 15° angle, 0.8 mm depth from the dura, 3 m/s velocity, and 100 ms dwell time. Sutures were administered as above.

### tMCAO

Mice (age P25-45) underwent focal ischemia-reperfusion with transient right middle cerebral artery occlusion (tMCAO) for 3 hours, or sham surgery, as detailed previously.^69–71^ Briefly, the right internal carotid artery (ICA) was dissected and a temporary ligature was tied using a strand of 6-0 suture at its origin. This ligature was retracted laterally and posteriorly to prevent retrograde blood flow. A second suture strand was looped around the ICA above the pterygopalatine artery and an arteriotomy was made proximal to the isolated ICA. A silicone coated 6-0 nylon filament from Doccol Corporation (Sharon, MA, USA) was inserted 6.5-7 mm to occlude the MCA and the second suture strand was tied off to secure the filament for the duration of occlusion. Injury was confirmed by severe left frontal/hindlimb paresis resulting in circling movements during the occlusion period. For reperfusion, each animal was anesthetized and all suture ties and the occluding filament were removed. Avitene Microfibrillar Collagen Hemostat (Warwick, RI, USA) was placed over the arteriotomy and the skin incision was closed. Sham animals were anesthetized, and the ICA was dissected, after which the skin incision was closed. At the time of reperfusion, the sham animals were once again anesthetized for 5 minutes, equivalent to the reperfusion procedure time for tMCAO animals.

### Marmoset stroke

Induction of focal stroke to the marmoset primary visual cortex (V1) was performed by vasoconstrictor-mediated vascular occlusion of the calcarine branch of the posterior cerebral artery (PCAca), as detailed previously.^72,73^ Briefly, following anesthesia (Alfaxalone 5mg/kg; maintained with inspired isoflurane 0.5-4%), a craniotomy and dural thinning was performed, followed by intracortical injections of endothelin-1 (ET-1; 0.1µl/30s pulse at 30s intervals, totaling ∼0.7 µL over 7 sites) proximal to the PCAca, which supplies operculum V1. The craniotomy was replaced, secured with tissue adhesive (Vetbond; 3M) and the skin sutured closed. Monkeys recovered for 7 days (n=2), 6 weeks (n=2) and 1 year (n=2; equal sex).

### Liposome injection

Clodronate liposomes or empty control liposomes (Encapsula Nanoscience) were administered intraperitoneally, beginning with injection 3 days prior to injury (200μL) followed by injection on the day of injury (100μL) and every 3 days subsequently (100μL) until harvest (14dpi).

### Antibody injection

ADWA11^74,75^ (generously provided by Dean Sheppard) or IgG1 isotype control (InVivoMab) was diluted in sterile DPBS to 10mg/kg and injected intraperitoneally on the day of injury (0dpi) and every week subsequently until harvest (7dpi, etc.).

### EdU injection

To measure proliferation, 5-ethynyl-2’-deoxyuridine (EdU, Thermo Scientific) was reconstituted at 5mg/mL in DPBS and injected at 50mg/kg i.p. Mice were injected either every other day throughout the injury timecourse or 2 hours prior to sacrifice.

### Tissue processing: imaging (mouse)

Following CO_2_ euthanasia, mice were transcardially perfused with 10mL of DPBS and 10mL of 4% PFA (Thermo Scientific). Skullcaps/brains were removed from bases. Skullcaps (meningeal imaging) or skullcaps/brains were fixed overnight (4% PFA, 4°C). Skullcaps/brains were washed (DPBS) and decalcified in 0.3M EDTA (VWR) (1 week, 4°C) followed by cryoprotection (30% sucrose). When removed from skullcaps, brains were cryoprotected directly after fixation. Brains were frozen in O.C.T. (Thermo Scientific) on dry ice and sliced to indicated thickness on a cryostat (Leica). Spinal cords processed similarly (decalcifying intact vertebra). For spatial transcriptomics, tissue processing was performed as above, without PFA perfusion/fixation. Brains were removed from skullcaps and directly frozen. For quantitative imaging of (blinded) brains, 2 14μm sections were collected per 100μm sliced, and sections representing a lesion’s maximal cross-sectional area were stained, imaged, and quantified. Lesion size outliers (diameter <25% of normal, representing technical errors during injury) were excluded prior to unblinding, resulting in a single exclusion from **Fig. 4k, Extended Data Fig. 9p**. Mice with hydrocephalus were also excluded.

### Tissue processing: imaging (marmoset)

For immunofluorescence, naïve controls (n=2; equal sex) and post-stroke marmosets were administered an overdose of pentobarbitone sodium (100mg/kg; intraperitoneal). Following apnea, animals were transcardially perfused with 0.1M heparinized PBS, followed by 4% paraformaldehyde (PFA) in PBS (0.1M). Brains were dissected, post-fixed and cryoprotected, as outlined previously.^72,73,76^ Following separation of the hemispheres, each hemisphere was bisected coronally at the start of the caudal pole of the diencephalon and frozen in liquid nitrogen at-40°C. Tissue was cryosectioned in the parasagittal plane at-20°C to obtain 40µm for free-floating sections stored in cyroprotectant solution (50% PBS 0.1M, 20% ethylene glycol, 30% glycerol).

### IHC

Slide-mounted thin sections were thawed, washed (DPBS), and blocked (1 hour, DPBS/0.4% TritonX-100/5% secondary-host serum). Samples were incubated in primary antibodies diluted in blocking solution (room temperature, 1 hour, or 4°C, overnight). Samples were washed (DPBS/0.05% TritonX-100, 5 minutes, 3 times) and incubated in secondary antibodies diluted 1:1000 in blocking solution (room temperature, 45-60 minutes). Samples were washed, mounted in DAPI Fluoromount-G (Thermo Scientific), and imaged. Proliferation was measured using the Click-iT EdU Alexa Fluor 647 Imaging Kit (Thermo Fisher), according to the manufacturer’s instructions (in between primary and secondary staining steps).

Medium-thickness sections (30-50μm) were blocked in 250μL DPBS/0.25% TritonX-100/5% secondary-host serum (1 hour, room temperature). Samples were subsequently incubated in primary antibody diluted in blocking solution (4°C, overnight), washed (DPBS/0.05% TritonX-100, 5 minutes, 4 times), and incubated in secondary antibodies diluted 1:500 in blocking solution (room temperature, 45-60 minutes). Samples were washed 3 times, mounted in DAPI Fluoromount-G, and imaged.

Meninges were blocked and stained before removal from skullcaps (block: 4°C, overnight, in 2mL DPBS/0.3% TritonX-100/5% FBS/0.5% BSA/0.05% NaN_3_). Samples were incubated in primary antibody diluted in 600μL of staining solution (DPBS/0.15% TritonX-100/7.5% FBS/0.75% BSA/0.075% NaN_3_; 4°C, 72 hours), washed (DPBS/0.15% TritonX-100, 4°C, 30 minutes, 3 times), and incubated in secondary antibodies diluted 1:400 in staining solution solution (4°C, 24 hours). Samples were washed and incubated in 10μg/mL DAPI in PBS (room temperature, 1 hour). Dural meninges were subsequently microdissected from skullcap, mounted in 50-100μL of Refractive Index Matching Solution (RIMS, DPBS/Histodenz [133.33g/100mL]/0.017% Tween-20/0.17% NaN_3_), and imaged.

Thick sections (100-200μm) were stained using the iDISCO protocol^77^ with 1-2 days of permeabilization, 1-2 days of blocking, 3 days of primary antibody, and 3 days of secondary antibody.

For marmoset imaging, free-floating sections comprising the infarct and peri-infarct regions were selected and washed in PBS (0.1M) and pre-blocked in a solution of 10% normal goat serum in PBS + 0.3% Triton X-100 (TX; Sigma) before incubation with primary antibodies overnight at 4°C. Sections were rinsed in 0.1% PBS-Tween and incubated with secondary antibodies (1 hour). After washes in PBS, sections were treated with DAPI, mounted in Fluoromount-G, and imaged.

### Imaging Antibodies

Primary antibodies used for murine imaging include chicken anti-GFP (Aves Labs GFP-1020, 1:200), rabbit anti-dsRed (Takara 632496, 1:300), chicken anti-GFAP (Invitrogen PA1-10004, 1:200), rat anti-GFAP (2.2B10, Invitrogen 13-0300, 1:200), rat anti-ER-TR7 (Novus Biologicals NB100-64932, 1:200), rabbit anti-*α*SMA (Abcam ab5694, 1:300) rat anti-CD31 (MEC13.3, Biolegend 102514, 1:200), goat anti-Desmin (GenWay Biotech GWB-EV0472, 1:200), goat anti-Decorin (Novus Biologicals AF1060, 1:200), rabbit anti-Collagen 6*α*1 (Novus Biologicals NB120-6588, 1:200), rat anti-Periostin (345613, Novus Biologicals MAB3548, 1:200), rat anti-ICAM1 (YN1/1.7.4, Biolegend 116110, 1:200), syrian hamster-anti-CD3*ε* (500A2, BD Biosciences 553238, 1:200), goat anti-S100A8 (R&D Systems AF3059, 1:200), chicken anti-NeuN (Millipore Sigma ABN91, 1:200), rabbit anti-Iba1 (Aif3, Fujifilm Wako 019-19741, 1:200-1:1000). For marmoset imaging, rabbit anti-COL6 (Abcam ab6588, 1:500) was used.

Secondary antibodies were used as necessary at protocol-specific concentrations, conjugated to A488, A555, AF594, and A647 (Life Technologies, Thermo-Fisher).

### Confocal and widefield microscopy

Confocal images (for thick sections and T-cell quantification) were imaged using a Nikon A1R laser scanning confocal including 405, 488, 561, and 650 laser lines for excitation and imaging with 16X/0.8 NA Plan Apo long working distance water immersion, 20X/0.95 NA XLUM PlanFl long working distance water immersion, or 60X/1.2 NA Plan Apo VC water immersion objectives. Z steps were acquired every 4μm. Widefield images (for thin sections, fibrosis quantification, and lesion size quantification) were imaged using a Zeiss Axio Imager.M2 widefield fluorescent microscope with a 10X/0.3 air objective. Marmoset images were acquired using the VS200 slide scanner (Olympus).

### Image analysis/quantification

z-stacks were rendered in 3D and quantitatively analyzed using Bitplane Imaris v9.8 software package (Andor Technology PLC, Belfast, N. Ireland). Individual cells (e.g., lymphocytes) were annotated using the Imaris surface function based on fluorescent CD4^cre^; Rosa26^Ai14^ signal when available, along with CD45, Iba1, and/or CD3e staining, DAPI staining, and size/morphological characteristics, with background signal in unrelated channels excluded. 3D reconstructions of fibrotic and glial scars were generated using the Imaris surface function based on fluorescent ER-TR7 or GFAP signal. 3D distances between lymphocytes and stromal/glial scars were calculated using the Imaris Distance Transform Matlab extension. MFI values of individual channels within a given distance of stromal/glial structures were determined using the Imaris Dilate Surface Matlab extension (the median value per image was recorded). Proportions of immune cells within each region were determined by manually tracing GFAP/ER-TR7 borders and filtering immune cell surfaces based on inclusion. All T-cell quantifications were performed within rectangular regions of a standard area centered on lesions. When applicable, fibroblasts were annotated similarly, using the Imaris surface function based on Col1a2^creER^; Rosa26^Ai14^ and Col1a1^GFP^ fluorescence, along with DAPI staining, size, and morphological characteristics. Similar surfacing thresholds for given cell types and structures were applied across slices and experiments, modifying individual thresholds when necessary to correct for varying tissue background.

Lesion sizes were calculated in Fiji (ImageJ version 1) by tracing ER-TR7 and GFAP borders. Fibroblast or myeloid cell coverage was determined by thresholding the relevant channel (e.g., ER-TR7), using a consistent threshold for each slice except for cases of exceptionally high tissue background. Coverage was calculated as thresholded area normalized to lesion area, and data was subsequently unblinded. For pooled data, equivalent thresholds were applied across experiments when possible, with exceptions made for varying tissue background.

### Tissue Processing: Visium

Tissue was collected from 1 resting mouse, 1 2dpi mouse, 2 7dpi mice, and 4 21dpi mice (including 2 Cthrc1^creER^; R26^DTA^ mice, discarded after initial clustering due to undetectable deletion; PT injury). Tissue was collected as above. 10μm slices were prepared via cryostat and directly mounted onto Tissue Optimization and Spatial Gene Expression slides (10X Genomics). Tissue optimization was carried out as per manufacturer’s recommendations, resulting in an optimal tissue permeabilization time of 12 minutes. Tissue mounted on the Spatial Gene Expression slide was processed as per manufacturer’s recommendations, imaged on a Leica Aperio Versa slide scanner at 20X, and transferred to the Gladstone Genomics Core for library preparation according to the manufacturer’s protocol. Samples were subsequently transferred to the UCSF Center for Advanced Technology for sequencing on the NovaSeq 6000 system.

### Tissue processing: nuclear isolation (murine)

Nuclear isolation was employed to overcome limitations of traditional flow cytometry and/or scRNAseq, including ECM interference with fibroblast isolation and dissociation signatures that disproportionately affect stromal and immune cells.^78^ For nuclear flow cytometry, tissue was collected from resting or 14dpi (PT injury) Pdgfr*α*^GFP^ mice. For single nuclear RNA sequencing experiment 1 (timecourse), tissue was collected from 2 wildtype mice per timepoint (1 male and 1 female) at rest, 2dpi, 7dpi, and 21dpi. 2 mice/timepoint with impaired TGF*β* signaling (Col1a2^creER^; Tgfbr2^flox^ and Cdh5^creER^; Tgfbr2^flox^) were harvested at 7/21dpi but were discarded after initial clustering and not analyzed separately due to insufficient yield. Alternatively, for single nuclear RNA sequencing experiment 2, tissue was collected from 2 WT mice, 2 Col1a2^creER^; Tgfbr2^flox^ mice, and 2 ADWA11-treated mice at 7 and 21dpi (1 male and 1 female per timepoint/condition).

Following CO_2_ euthanasia, mice were transcardially perfused (10mL DPBS) and decapitated. Brains were removed from skullcaps and placed in iMED+ (15mM HEPES [Fisher] and 0.6% glucose in HBSS with phenol red).^79^ For nuclear flow cytometry and single nuclear RNAseq experiment 1 (timecourse), dura, lesion, and perilesional cortex were microdissected and processed separately. For RNAseq experiment 2 (WT, cKO, and ADWA11), only lesions were dissected. Microdissection involved meningeal/skullcap separation, removal of subcortical structures, and separation of lesions from skullcaps (lesions often separate from cortex during initial dissection but can be micro-dissected as necessary). For snRNAseq, tissue from male and female mice within experimental conditions was combined.

Tissue was processed using ST-based buffer protocol,^78^ with the following modifications: initial centrifugation was performed at 500g for 10 minutes. After lysis/initial centrifugation, nuclei were resuspended in 1mL ST buffer (nuclear flow cytometry, RNAseq experiment 2) or PBS/1% BSA/0.2UμL Protector RNAse inhibitor (Roche) (RNAseq experiment 1), filtered through 35μm cell strainers, and subsequently processed as below.

For nuclear flow cytometry and single nuclear RNAseq experiment 2, nuclei were centrifuged for 5 minutes at 500g, resuspended in FANS buffer (DPBS/1% BSA/0.1mM EDTA^80^) with 2μg/μL DAPI and 0.2U/μL RNase inhibitor (snRNAseq), and stained (nuclear flow) or sorted (snRNAseq).

For single nuclear RNAseq experiment 1, cell counts were performed after initial centrifugation (NucleoCounter, Chemometic), and a maximum of 2×10^6^ nuclei were multiplexed using CellPlex Multiplexing technology (10X Genomics) according to the manufacturer’s instructions (using protocol 1 for nuclear multiplexing, with only one wash after multiplexing to increase yield). Nuclei were resuspended in FANS buffer with 0.2U/μL RNase inhibitor and 2μg/μL DAPI and nuclear concentrations were determined. Immediately before sorting, multiplexed microanatomical regions (including lesion, parenchyma, and dural meninges) from individual mice were combined at desired ratios (75% lesion, 25% parenchyma). Nuclei were sorted (forward and side scatter and DAPI) into pre-coated tubes containing 200μL PBS/1% BSA/0.2U/μL RNase inhibitor (BDFACSAria II sorting system, 100μm nozzle size, 4 way-purity sort mode). Resting dural fibroblasts were enriched using Col1a2^creER^; Rosa26^Sun1GFP^ and resting nuclei were combined after sorting (50% GFP^+^ meninges, 25% GFP^-^ meninges, 25% parenchyma).

After sorting/pooling (snRNAseq), samples were centrifuged (500g, 15 minutes), and supernatant was removed to leave a minimum final volume of 45μL with a maximum of 16,500 nuclei. Final nuclear concentrations were acquired, and samples were transferred to the UCSF Genomics CoLab for library preparation. Up to 1.6 x 10^4^ nuclei were loaded onto the Chromium Controller (10X Genomics). Chromium Single Cell 3’ v3.1 reagents were used for library preparation according to the manufacturer’s protocol. Libraries were transferred to the UCSF Institute for Human Genetics for sequencing on the NovaSeq 6000 system.

### Tissue processing: nuclear isolation (marmoset)

For single-nuclei RNA sequencing (snRNAseq), naïve control marmosets (n=3; 1 female, 2 male; median age 4Y) were administered an overdose of pentobarbitone sodium (100 mg.kg^-1^; intraperitoneal). Following apnea, frontal lobes were recovered and dissected under aseptic conditions in sterile ice-cold phosphate buffered saline (PBS; 0.1M; pH 7.2). Tissues were and snap frozen in isopentane chilled in liquid nitrogen. The procedures/ dissections were performed in chilled RNAase-free PBS with RNase-free sterilized instruments under RNase-free conditions. Approximate time from apnea to snap-freezing ranged from 20-30 minutes. All 6 samples passed QC. Nuclear isolation was performed as described previously,^76^ involving pulverization in liquid nitrogen, lysis in lysis buffer, dounce homogenization, and gradient purification. After isolation, nuclei were counted and diluted to 1 million/mL with sample-run buffer (0.1% BSA, RNAse inhibitor [80U/mL], 1mM DTT in DPBS). snRNAseq was performed on the 10x Genomics Chromium System. Cellranger commercial software was utilized to conduct initial data processes including sequence alignment to the marmoset genome (CalJac3).

### Tissue processing: single cell isolation (murine)

Single cell suspensions were prepared from tissues including brain, spinal cord, meninges, blood, and spleen. Immediately following CO_2_ euthanasia, spleens were removed into RPMI/10% FBS and peripheral blood was collected through the right ventricle into heparin tubes. Mice were subsequently transcardially perfused through the left ventricle with 10mL of DPBS, decapitated, and brains were carefully removed from skullcaps and placed in iMED+ as above.^79^ For select experiments, spinal cords were carefully dissected from vertebra. Cortex, lesion, and meninges were dissected as above. Brain was weighed and subsequently homogenized in iMED+ homogenized using a 2mL glass tissue grinder (VWR; 6 plunges, followed by filtration through a 70μm filter, addition of 2mL iMED+, and 6 more plunges). Filtered suspensions were centrifuged at 220g for 10 minutes and resuspended in 5mL of 22% Percoll (GE Healthcare) in Myelin Gradient Buffer (5.6mM NaH_2_PO_4_•H_2_O, 20mM Na_2_HPO_4_•2H_2_O, 140mM NaCl, 5.4mM KCl, 11mM glucose in H_2_O).^79^ 1mL of PBS was layered on top of Percoll. Samples were centrifuged at 950g for 20 minutes at 4°C with no break to separate myelin and resuspended in FACS buffer. Dissected meninges were incubated in digestion medium (RPMI/10% FBS/80μg/mL DNase I/40μg/mL Liberase TM (Roche)). Tissue was subsequently mashed through 70μm filters, followed by centrifugation and resuspension in FACS buffer. Spleens were prepared by mashing tissue through 70μm filters without tissue digestion, followed by centrifugation. Red blood cells were lysed for 2 minutes using 1X Pharm-Lyse and the remaining cell pellet were resuspended in FACS buffer. Blood samples were centrifuged for 5 minutes at 500g. Pellets were resuspended in 1X Pharm-Lyse 5 minutes at room temperature, followed by centrifugation and final suspension in FACS buffer.

### Flow cytometry

Resuspended samples were stained in 96-well V-bottom plates. Surface staining was performed at 4°C for 45 minutes in 50μL staining volume. For experiments involving intra-cellular staining, cells were fixed and permeabilized using Foxp3 Transcription Factor Staining Buffer Set (eBioscience) followed by staining at 4°C for 1 hour in 50μL staining volume. All samples were acquired on a BD LSRII Fortessa Dual or a BD FACSAria II for cell sorting. Live cells or nuclei were gated based on their forward and side scatter followed by Zombie NIR fixable (Biolegend 423106), Fixable Viability Dye eF780 (eBioScience 65086514), Draq7 (Biolegend 424001), or DAPI (40,6-diamidine-20-phenylindole dihydrochloride; Millipore Sigma D9542-10MG) exclusion (or inclusion for nuclei). Lineages were subsequently identified as follows:

Oligodendrocyte-lineage nuclei were identified as DAPI^+^, Pdgfr*α*^GFP-hi^, Olig2^+^. Fibroblast nuclei were identified as DAPI^+^, Pdgfr*α*^GFP-int^ or DAPI^+^, Col1a2^creER^; Rosa26^Sun1GFP+^ (snRNAseq experiment 1, resting dural meninges). Bulk nuclei were identified as DAPI^+^. Global lymphocytes were defined as CD45^+^, Thy1^+^. T cells were identified as CD45^+^, CD11b^-^, CD19^-^, NK1.1^-^, CD3*ε*^+^, CD4^+^ (CD4 T cells), or CD8*α*^+^ (CD8 T cells), and were further defined as CD44^+^, CD69^+^ (resident memory T cells, TRM), CD62L^+^ (naïve T cells), CD62L^-^, CD44^+^ (activated T cells), or CTV^diluted^ (proliferating T cells). Additionally, CD4 T cells were defined as T-bet^+^ (Th1 T cells), Gata3^+^ (Th2 T cells), or ROR*γ*t^+^ (Th17 T cells). Neutrophils were defined as CD45^+^, CD11b^+^, Ly6G^+^ (and optionally Thy1^-^, CD19^-^, NK1.1^-^). Monocytes were defined as CD45^+^, CD11b^+^, Ly6G^-^, Ly6C^+^ (and optionally Thy1^-^, CD19^-^, NK1.1^-^, Siglec F^-^). Microglia were defined as CD45^int^, CD11b^+^. Macrophages were defined as CD45^+^, Ly6G^-^, Ly6G^-^, CD64^+^ (optionally MERTK^+^) and were further defined as Scar Associated Macrophages (SAM, CD9^+^ and TREM2^+^). cDCs were identified as CD45^+^, Ly6G^-^, Ly6C^-^, CD64^-^, MHCII^+^, CD11c^+^, and were further defined as cDC1s (CD11b^lo^, optionally SIRP*α*^-^) or cDC2s (CD11b^hi^, optionally SIRP*α*^+^). B cells were defined as CD45^+^, Thy1^-^, CD19^+^. Eosinophils were defined as CD45^+^, Thy1^-^, CD19^-^, NK1.1^-^, Ly6G^-^, CD11b^+^, Siglec F^+^. Populations were back-gated to verify purity and gating.

Data were analyzed using FlowJo software (TreeStar, USA) and compiled using Prism (Graphpad Software). Cell counts were performed using flow cytometry counting beads (CountBright Absolute; Life Technologies) per manufacturer’s instructions.

### Flow cytometry Antibodies

Antibodies used for flow cytometry include rabbit anti-Olig2 (Thermo P21954), anti CD45 (30-F11, BD Biosciences 564279), anti-CD90.2 (Thy1, 53-2.1, Biolegend 140327, BD Biosciences 553004), anti-CD11b (M1/70, Biolegend 101224 or BD Biosciences 563015), anti-CD19 (6D5, Biolegend 115554), anti-NK1.1 (PK136, Biolegend 108736), anti-CD3*ε*(17A2, Biolegend 100216), anti-CD4 (RM4-5, Biolegend 100557, or GK1.5, BD Biosciences 563050), anti-CD8*α* (53-6.7, Biolegend 100750), anti-CD44 (IM7, Biolegend 103030), anti-CD69 (H1.2F3, Biolegend 104505), anti-CD62L (MEL-14, Biolegend 104407), anti-Tbet (4B10, Biolegend 25-5825-80), anti-Gata3 (TWAJ, eBioscience 12-9966-41), anti-ROR*γ*t (B2D, eBioscience 17-6981-82), anti-Ly6G (1A8, Biolegend 127624), anti-Ly6C (HK1.4, Biolegend 128011 or 128035), anti-CD64 (X-54-5/7.1, Biolegend 139323 or BD Biosciences 558539), anti-MERTK (DS5MMER, eBioscience 46-5751-80), anti-CD9 (KMC8, BD Biosciences 564235), anti-TREM2 (237920, R&D systems FAB17291A), anti-I-A/I-E (MHCII, M5/114.15.2, BD Biosciences 748845), anti-CD11c (N418, Biolegend 117339 or 117318), anti-CD172a (SIRP*α*, P84, eBioscience 12-1721-80), and anti-Siglec-F (E50-2440, BD Biosciences 740956).

### Ex vivo coculture

For *ex vivo* coculture experiments, lesions and contralateral cortex were dissected as described above and divided into halves or equivalently sized sections (contralateral cortex). Tissue was added to 96-well round bottom plates in 100μL of R10 (RPMI supplemented with 10% FBS, 1% penicillin/streptomycin, 1X Glutamax (Thermo Scientific), and 55μM b-mercaptoethanol). T-cells were isolated from spleens and cervical/inguinal lymph nodes using EasySep Magnetic Bead negative selection (Stem Cell), according to the manufacturer’s instructions (using 31.5μL of vortexed selection beads). A maximum of 1×10^7^ T-cells were labeled with 5mM CellTrace Violet in PBS (Thermo Scientific) for 10 minutes, followed by washing with R10, centrifugation, and resuspension at 1×10^6^ cells/mL. 100μL of suspension (containing 100,000 T-cells) was subsequently plated with lesions, control tissue, or alone. As a positive control, anti-CD3/CD28 T-cell activating DynaBeads (Thermo scientific) were magnetically washed in 1mL DPBS and added to T-cells at a 1:1 ratio. Plates were incubated at 37°C, 5% CO_2_ for 72 hours. After coculture completion, wells were mixed by pipetting and cells were transferred to a V-bottom plate, and blocked, stained, and analyzed as above. CTV staining was used to identify plated (vs. lesion-resident) T cells. **Fig. 3m** and **Extended Data Fig. 7m-p** include pooled control data from experiments where wildtype lesions were cultured with diphtheria toxin (100ng/mL; any lesions expressing Rosa26^DTR^ were excluded), empty liposomes, or IgG1 isotype control (BioXCell). For direct comparisons between 7dpi and 21dpi lesions, media refeeding was performed daily (replacing 100µL, 2x) to mitigate rapid media acidification by 7dpi lesions.

### Vascular permeability analysis

Vascular permeability was measured using Evans Blue extravasation.^81^ Briefly, Evans Blue (1% in saline) was injected i.p. (8mL/kg) 3 hours prior to sacrifice. Following euthanasia, mice were transcardially perfused (10mL DPBS), and mice that did not show appropriate liver clearing were excluded. Brains were dissected as above, whole hemispheres were weighed and added to 500μL formamide, and tissue was incubated for 48 hours to extract Evans Blue. Evans Blue fluorescence was measured on a SpectraMax microplate reader (excitation 620nm, emission 680nm). A standard curve was generated and used to calculate extravasated mass, which was subsequently normalized to tissue weight.

### Visium data processing

Sequencing data were aligned to mouse genome mm10 with SpaceRanger version 2.0.0 (10x Genomics). Data was processed using the Seurat R package, version 4.2.1.^82^ Individual capture areas were processed using Seurat’s *SCTransform* function to normalize data, select variable features for dimensionality reduction, and scale data. Capture areas were subsequently merged for further analysis. Principle components were calculated using Seurat’s *RunPCA* function, followed by graph-based clustering using Seurat’s *FindNeighbors* (dims =1:30) and *FindClusters* (res = 0.8) functions and 2D visualization using Seurat’s *RunUMAP* function (dims = 1:30). Feature and spatial feature plots, violin plots, and UMAP plots were generated using Seurat. We used *FindAllMarkers* to identify markers for each cluster (test.use = MAST, min.pct = 0.05, logfc.threshold = 0.2) and generated resulting dot plots using Seurat. Fibroblast-containing clusters were identified via expression of *Col1a1* and further investigated using feature plots and spatial feature plots. One large cluster (cluster 10) was subclustered using Seurat’s *FindSubCluster* function (res = 0.5); subcluster 10_0 was identified as a 7dpi fibroblast-enriched cluster and selected for further analysis based on expression of *Pdgfra*. Cluster 8 was identified as a 21dpi fibroblast-enriched cluster.

Analyzing WT tissue, we used Seurat’s *FindMarkers* function (min.pct = 0.05, logfc.threshold = 0.2) to determine markers for cluster 10_0 and 8 and used the EnhancedVolcano R package to visualize relevant DEGs (*EnhancedVolcano* function, minfc = 1.15, alpha = 0.05).^83^ To calculate fibroblast TGF*β* scores, we utilized a previously generated bulk sequencing dataset of primary lung fibroblasts treated with TGF*β* (1ng/mL) or PBS for 48 hours.^84^ TGF*β*-upregulated genes were identified from normalized data (log2FC > 0, q < 0.05). Module scores were generated using Seurat’s *AddModuleScore* function. To calculate proliferation scores, we utilized a proliferation signature composed of 23 genes (*Ccnb1, Ccne1, Ccnd1, E2f1, Tfdp1, Cdkn2b, Cdkn1a, Plk4, Wee1, Aurkb, Bub1, Chek1, Prim1, Top2a, Cks1, Rad51l1, Shc, Racgap1, Cbx1, Mki67, Mybl2, Bub1, Plk1*).^85,86^

### Murine snRNAseq data processing

Sequencing data were aligned to mouse genome mm10 with CellRanger version 7.1.0 (snRNAseq experiment 1; timecourse) or version 7.2.0 (snRNAseq experiment 2; WT, cKO, and ADWA11) (10x Genomics). Data was processed using the Seurat R package, version 4.2.1 (experiment 1) or 5.0.1 (experiment 2). We excluded cells with high mitochondrial gene expression and low or high UMI and feature counts, using the bottom and top 2.5 percentiles as our cutoff. We used Seurat’s *SCTransform* function to normalize data, select variable features for dimensionality reduction (with the removal of certain sex-related and mitochondrial genes), and scale data. Principle components were calculated using Seurat’s *RunPCA* function, followed by graph-based clustering using Seurat’s *FindNeighbors* (dims =1:30) and *FindClusters* (res = 0.5) functions and 2D visualization using Seurat’s *RunUMAP* function (dims = 1:30). We used *FindAllMarkers* (test.use = MAST, experiment 1) or *RunPrestoAll*^87^ (experiment 2) to identify markers for each cluster (min.pct = 0.05, logfc.threshold = 0.2) and annotated clusters using select marker genes. We identified and excluded 2 clusters made up of likely doublets based on gene expression and excluded 1 sample due to insufficient nuclear yield (experiment 1). After visualizing resultant clusters on UMAPs and dot plots using Seurat, we subset the data to wildtype cells for downstream analysis. Additional feature plots, UMAP plots and dot plots, and heatmaps were generated using Seurat. Joint densities for gene combinations (including published meningeal layer signatures^36^) were visualized using the NebulosaPlot R package (*plot_density* function)^88^ and modified using the ScCustomize R package (*Plot_Density_Custom* function).^89^

Fibroblasts and immune cells were subset and reclustered as above (res = 0.5 [experiment 1] or 0.2 [experiment 2] for fibroblasts, res = 0.15 [experiment 1] or 0.25 [experiment 2] for immune cells) and DEGs were recalculated. Resting dural meningeal fibroblasts were subsequently removed (by microanatomical metadata and cluster) for downstream analysis; pericytes were similarly removed for **Fig. 2e** and **Fig. 3b**. For experiment 2, clusters were annotated by expression of previously defined CNS fibroblast or myeloid cell “signatures” generated from experiment 1 DEG marker lists (visualized using *AddModuleScore* and violin plots). Dural fibroblasts were subclustered (*FindSubCluster* function) to identify lymphocyte-interactive and dural subsets. Relative abundance across time was calculated for each fibroblast and macrophage subcluster after normalizing for timepoint or condition sample size (i.e., total # nuclei). Gene ontology analysis was performed using the ClusterProfiler R package for gene set testing (*enrichGO* function).^90^ Ligand-receptor and ligand-signaling-network interactions underlying myofibroblast emergence were interrogated using the NicheNetR R package.^91^ Beginning with the global Seurat object, fibroblasts were treated as “receiver” cells, with resting fibroblasts as the “reference condition” and 7dpi myofibroblasts as the “condition of interest.” Potential “sender” cells were any cells present at rest, 2, or 7dpi. Integration of single nuclear and Visium data was performed using the SpaceXR package^92^ (to deconvolute individual spots) and using Seurat’s *AddModuleScore* function (to score each Visium cluster with marker sets for single nuclear fibroblast clusters). Multi-gene scores were calculated as follows and generated using *AddModuleScore.* Profibrotic “SAM scores” were generated as published previously, using a combination of 6 genes (*Trem2*, *Cd9*, *Spp1*, *Gpnmb*, *Fabp5*, and *Cd63*).^26^ Disease associated microglia (“DAM”) scores were generated using the top 30 published markers specific to DAM (p < 0.001, ranked by logFC).^33^ Dysmaturity scores were generated using genes significantly upregulated in microglia from Emx1^cre^; Itgb8^flox^ mice (relative to controls; bulk sequencing data accessed via the GEO and analyzed with originally described parameters).^39^ Macrophage-fibroblast ligand-receptor interactions were interrogated and visualized using CellPhoneDB (Version 4, statistical method).^93^ Pseudotime trajectories for myeloid cells were generated using the monocle3 R package v1.3.4 (*learn_graph* and *order_cells* functions), with homeostatic microglia and monocytes chosen as 2 possible roots.

### Marmoset snRNAseq data processing

Processed data from 7dpi marmosets (ET1-induced stroke)^76^ and resting marmosets (unpublished) was generously provided by the J. Bourne lab. Fibroblasts from both datasets were identified via COL1A1 expression, subset, and integrated using the Seurat *FindIntegrationAnchors* and *IntegrateData* functions (dims = 1:30). The integrated data was subsequently rescaled, PCAs were calculated, clusters were identified, and marker genes were selected as above. UMAP plots, barplots, and heatmaps of selected fibrosis-related genes were generated using Seurat.

### Human TBI snRNAseq data processing

Human TBI snRNAseq data was accessed via the GEO.^94^ Datasets representing individual patients were merged, and quality control was performed on mitochondrial expression, UMI counts, and gene counts, as above. We used Seurat’s *SCTransform* function to normalize data, select variable features for dimensionality reduction (with the removal of certain sex-related and mitochondrial genes), and scale data. Principle components were calculated using Seurat’s *RunPCA* function, followed by batch correction using Harmony,^95^ graph-based clustering, and cluster identification as above (res = 0.1). After cluster annotation, fibroblasts were identified via COL1A1 expression and subset. Heatmaps and violin plots were generated using Seurat.

### Human GBM scRNASeq data processing

Human GBM data was generously provided by the M. Aghi lab.^16^ QC was performed in accordance with the original publication (excluding cells with mt.percent > 20% and fewer than 200 or more than 20,000 UMIs). We used Seurat’s *SCTransform* function to normalize data, select variable features for dimensionality reduction (with the removal of certain sex-related and mitochondrial genes), and scale data. Principle components were calculated using Seurat’s *RunPCA* function, followed by graph-based clustering and cluster identification as above (res = 0.1). After cluster annotation, fibroblasts were identified via COL1A1 expression, subset, and reclustered as above (res = 0.6). UMAP plots, feature plots, and heatmaps were generated using Seurat.

### Statistical analysis

All data were analyzed by comparison of means using unpaired (unless otherwise noted) two-tailed Student’s t-tests; for multiple comparisons, one-way or two-way ANOVA (with Tukey or Sidak’s post hoc test) were used as appropriate (Prism, GraphPad Software, La Jolla, CA). ns = not significant, *p<0.05, **p<0.01, ***p<0.001, ****p<0.0001. Figures display means ± standard deviation (SD) unless otherwise noted. When possible, results from independent experiments were pooled. All data points reflect individual biological mouse replicates (flow analysis) or individual tissue slices analyzed (imaging), unless otherwise noted. For imaging experiments, all analysis was performed at least three mice from at least two independent experiments, with two or more fields analyzed per mouse.

### Data availability

Spatial and single nuclear transcriptomic data generated in this paper are deposited in Gene Expression Omnibus (GEO) (accession number will be released at the time of publication). Accession codes for 7dpi Marmoset data (Boghdadi et al.^76^), human TBI data (Garza et al.^94^), and human GBM data (Jain et al.^16^) can be found in the referenced studies.

### Code availability

All original code generated to analyze RNA sequencing data is available at GitHub (“https://github.com/newingcrystal/CNS_Fibroblasts”).

## Acknowledgments

We thank Dr. Richard Locksley for his constructive feedback on this manuscript, as well as the UCSF Parnassus Flow Core RRID:SCR_018206 and DRC Center Grant NIH P30 DK063720, the UCSF Biological Imaging and Development Core (BIDC), the UCSF Genomics CoLab, the UCSF Institute for Human Genetics, the Gladstone Histology & Light Microscopy Core (HLMC), the Gladstone Genomics Core, and the Monash Histology Platform for instruments and services. T.T. is supported by the NHLBI (NIH K99/R00HL155786). L.T. and J.A.B. are supported by the National Health and Medical Research Council (NHMRC; Australia; APP20140228) and The Yulgilbar Foundation Fund to JAB. The Australian Regenerative Medicine Institute is supported by the State Government of Victoria and the Australian Government. J.T.P. is supported by the NINDS (NIH R01NS096369). F.F.G. is supported by the NINDS (NIH R01NS107039). D.S. is supported by the NHLBI (NIH R01HL142568). T.D.A. is supported by the NINDS (NIH R01NS119615-01). J.T.P., A.B.M., and A.V.M. are supported by the NINDS (NIH R01NS126765), UCSF PBBR grant, and UCSF Dept. of Lab Medicine discretionary funds; A.B.M. is additionally supported by the NIAID (NIH R01AI162806).

## Author contributions

Conceptualization, NAEC, TDA, AVM, ABM; methodology, NAEC, NMM, SN, DS, TDA, ABM; investigation, NAEC, NMM, AAC, EDM, SEC, LT, AL, TT, AK, RP, GLM, SN, AC, HP, SJ, MKA, JAB, JTP, FFG, DS; resources, AVM, TDA, ABM; data curation, NAEC, NM; writing, NAEC, TDA, AVM, and ABM; supervision, TDA, ABM.

## Supplementary information

Supplementary Information is available for this paper.

## Lead contact

Further information and requests for resources and reagents should be directed to and will be fulfilled by the Lead Contact, Ari B. Molofsky.

---

**Extended Data Fig. 1:**
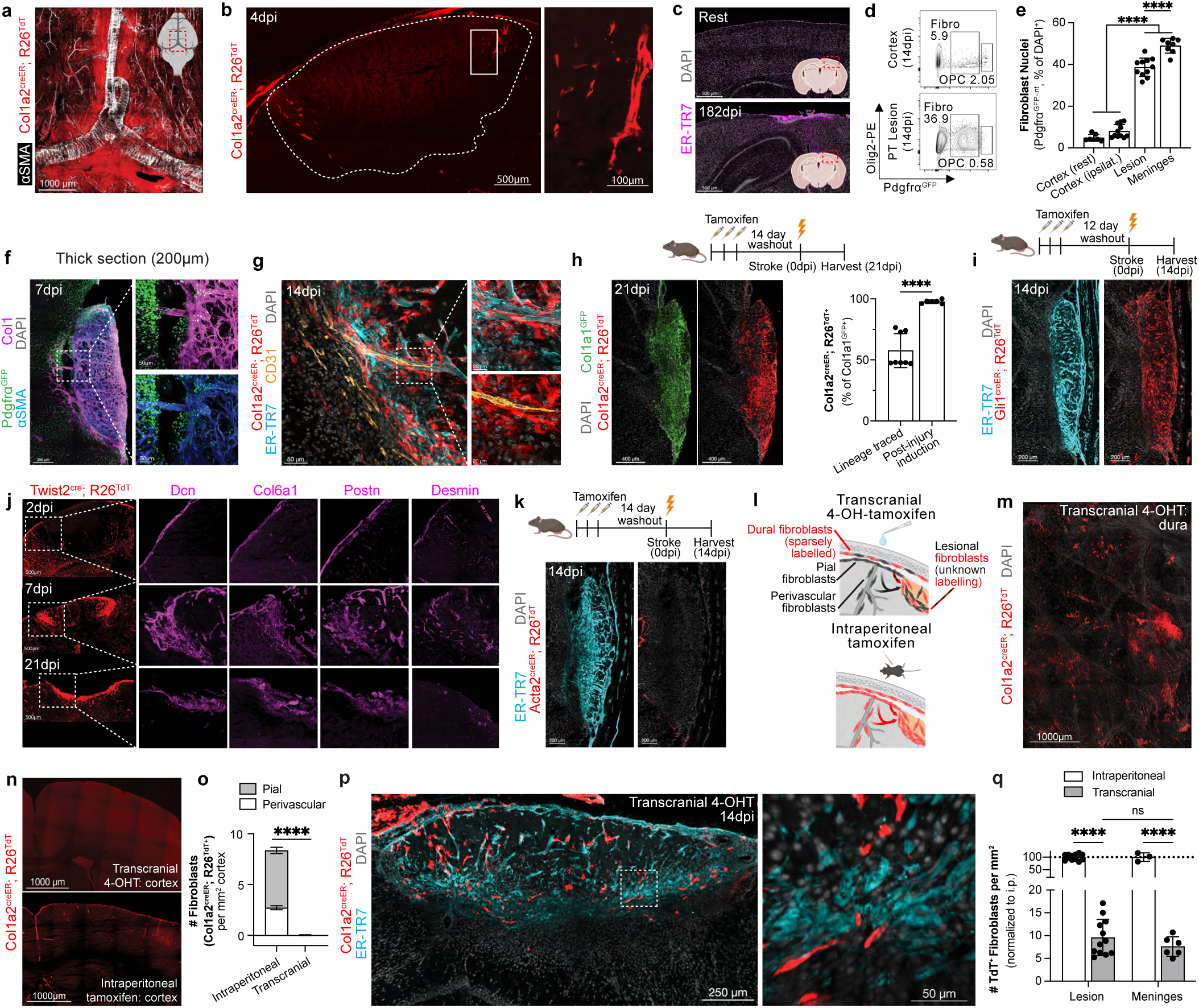
Additional characterization of the CNS fibroblast response to PT injury, related to Fig. 1. **a** Homeostatic dural meningeal fibroblasts (Col1a2^creER^; Rosa26^TdT+^; whole-mounted dura, tamoxifen days ^-^11-^-^7 before harvest). **b**, Fibroblast expansion near lesion borders (white dotted line) by 4dpi (tamoxifen days ^-^14-^-^12, ^-^7, 0-2dpi). **c**, Chronic cortical and subcortical/hippocampal contraction. **d-e**, Flow plot (**d**) and quantification (**e**) of fibroblast nuclei in Pdgfra^GFP^ mice (marking fibroblasts and oligodendrocyte precursor cells [OPC]). n=7 (resting cortex), 11 (ipsilateral cortex/lesion, 14dpi), or 8 mice (meninges, resting or 14dpi). **f-g**, Vascular remodeling after PT injury, with Pdgfra^GFP+^ nuclei (**f**) or fibroblasts (**g**) visualized in perivascular spaces. **h**, 21dpi fibroblasts (Col1a1^GFP+^) with a fibroblast origin (Col1a2^creER^; Rosa26^TdT+^). Tamoxifen days ^-^16-^-^14 (before injury, “lineage-traced”) or 7-9dpi (“post-injury induction”). n=8 (lineage-traced) or 6 (post-injury) slices (2 slices/mouse). **i**, Lineage traced fibroblasts in Gli1^creER^; Rosa26^TdT^ mice (preferentially recombining “universal” fibroblast subsets in multiple tissues). Tamoxifen days ^-^14-^-^12 before injury. **j**, Time course showing stromal/fibroblast (Twist2^cre^; Rosa26^TdT+^) expression of fibroblast markers (Dcn, Col6a1, Postn) and pericyte marker (Desmin). **k**, Lack of lineage traced fibroblasts in Acta2^creER^; Rosa26^TdT^ mice (TdT^+^ smooth muscle visible outside lesion). Tamoxifen days ^-^16-^-^14 before injury. **l-o,** Schematic (**l**) showing transcranial 4-hydroxy-tamoxifen (4-OHT) induction (left) and intraperitoneal tamoxifen induction (right). Transcranial 4-OHT sparsely recombines dural (**m**) but not pial or perivascular fibroblasts (**n-o**, shown 2-10 days after induction). n=5 (transcranial) or 3 (intraperitoneal) mice (**o**). **pq**, Image (**p**) and quantification (**q**) of lesional fibroblasts recombined via transcranial 4-OHT. n=12 slices from 6 mice (lesion, 2 slices/mouse), 6 mice (meninges/transcranial), or 3 mice (meninges/intraperitoneal). Graphs show mean±s.d. ns=not significant, ****p<0.0001; one-way ANOVA, Tukey post-test (**e**); Student’s T-test (**h**,**o**); two-way ANOVA, Sidak’s post-test (**q**). 14μm (**b**,**c**,**g-k**,**p**,**q**), 200μm (**f**), or 40μm (**j** [desmin], **n,o**) slices; images represent two or more mice.

**Extended Data Fig. 2:**
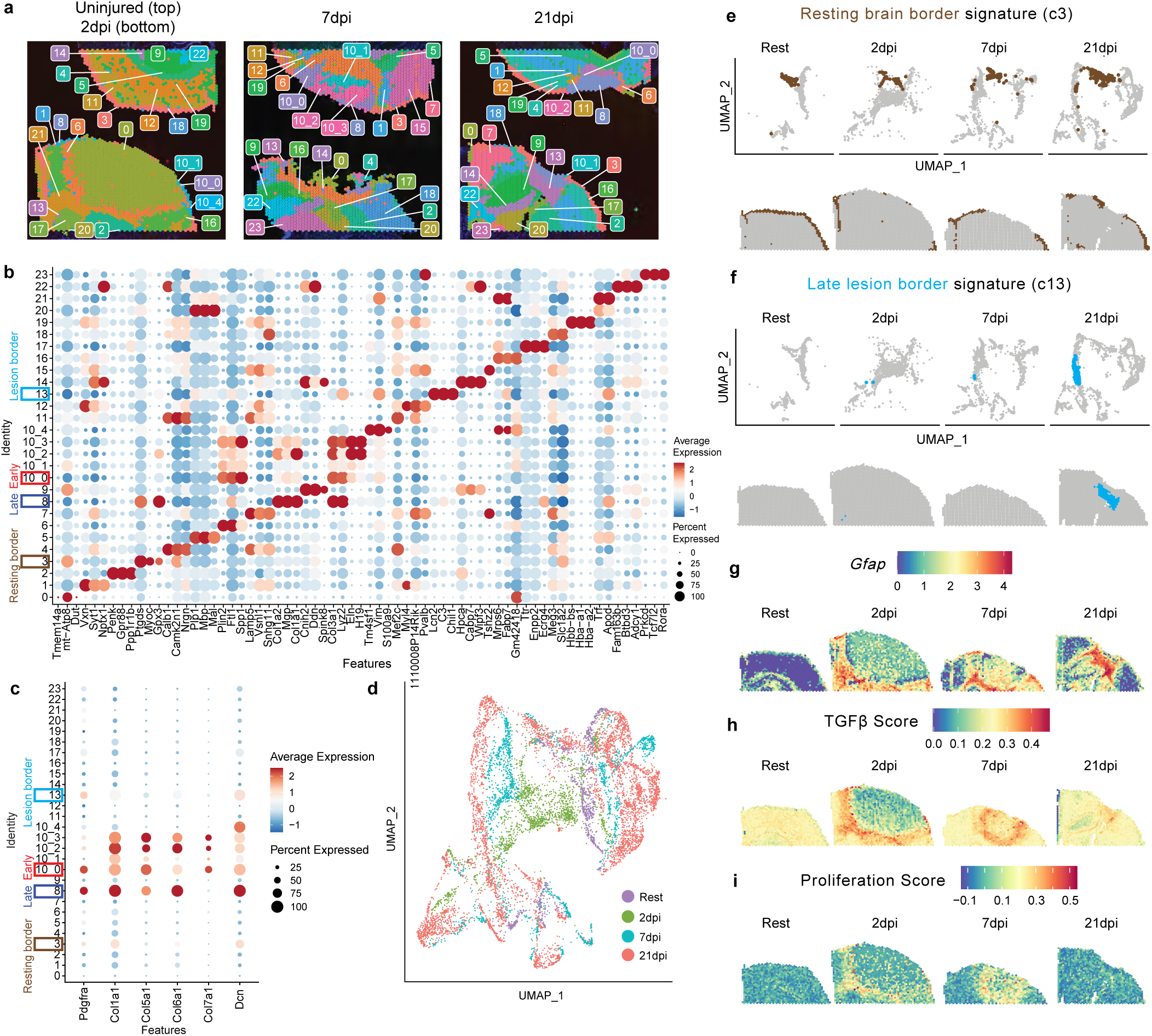
Spatial transcriptomics characterization of PT injury, related to Fig. 1. **a** Spatial transcriptomics plot showing all spot-based clusters. Slices shown in original Visium slide configuration with two slices per capture area: rest (top left), 2dpi (bottom left), 7dpi (middle, 2 biological replicates), and 21dpi (right, 2 biological replicates). **b-c**, Dot plots showing marker genes (**b**) or fibroblast-associated genes (**c**) for all spot-based clusters. c3 (“resting brain border”), c8 (“late fibroblast”), c10_0 (“early fibroblast”), and c13 (“late lesion border”) are highlighted (colored boxes). c10_0 was selected from c10 for further analysis based on high *Pdgfra* expression. **d**, UMAP showing spot-based clusters colored by timepoint, highlighting divergence among fibroblast-containing clusters. **e-f**, UMAP and spatial plots showing distribution of “resting brain border” fibroblast signature (**e**, cluster 3, brown) and “late lesion border” fibroblast signature (**f**, cluster 13, cyan). **g-i**, Spatial plots showing astrocyte-containing spots (**g**, *Gfap*^+^) and distribution of “fibroblast TGFb score” (**h**) or “proliferation score” (**i**).

**Extended Data Fig. 3:**
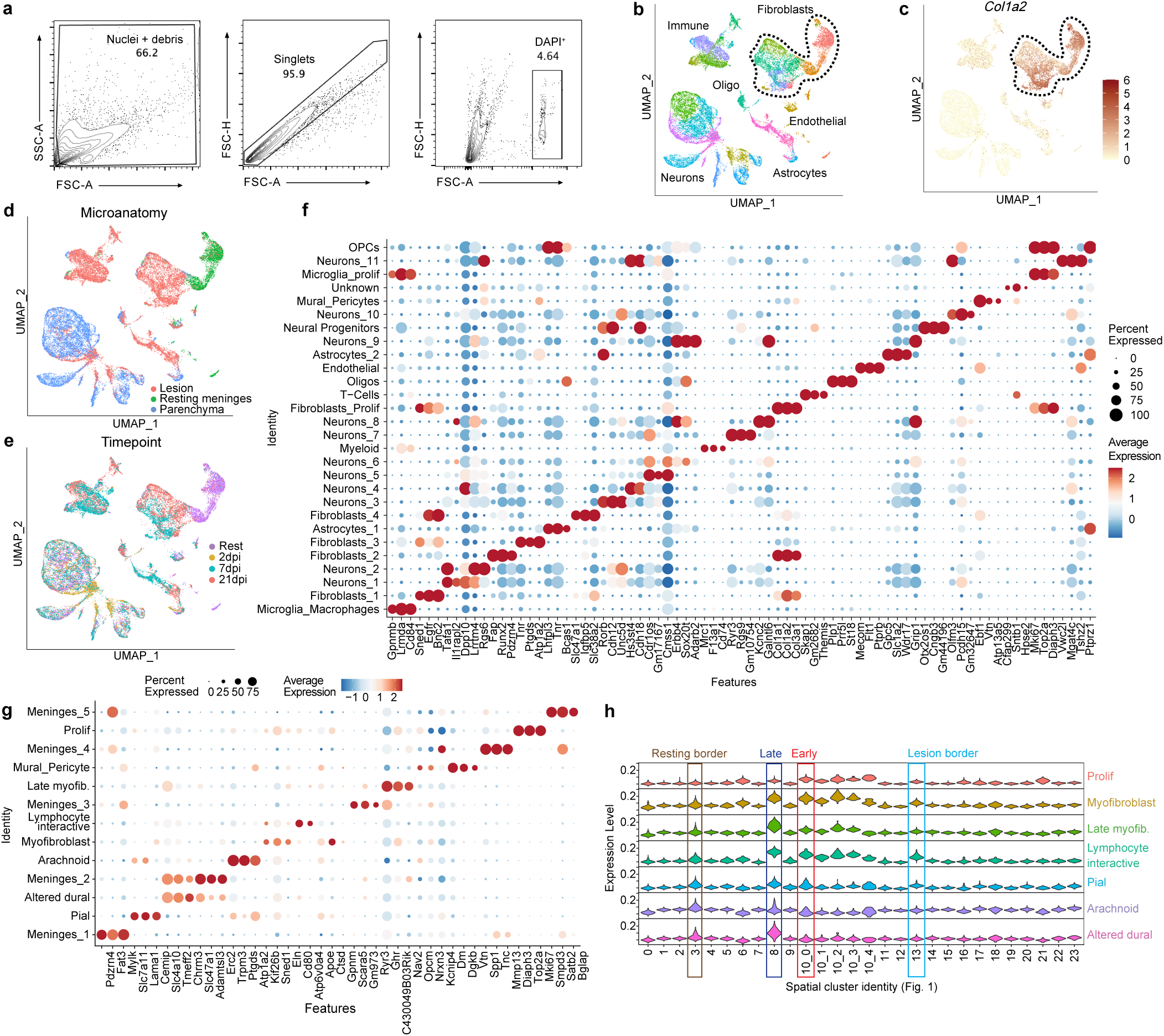
Additional transcriptomic characterization of fibroblasts in murine PT injury, related to Fig. 2. **a** Gating strategy for sorting DAPI^+^ nuclei for snRNAseq. **b-c**, Global UMAP (**b**) and *Col1a2* feature plot (**c**) showing fibroblast clusters (black dotted line) along with immune, oligodendrocyte lineage, endothelial, astrocyte, and neuron clusters. **d-e**, Global UMAPs showing barcode-based microanatomical metadata (**d**) or timepoint metadata (**e**). Neurons show parenchymal origin and broad temporal distribution, dural fibroblasts (dissected only at rest) show meningeal origin and resting distribution, and lesional fibroblasts, immune cells, and glial cell subsets show lesional origin and temporal heterogeneity. **f-g**, Dot plots showing marker genes for all cellular clusters (**f**) or all fibroblast subclusters (**g**). **h**, Violin plots mapping each lesional fibroblast cluster identity (via marker-DEGs) onto spatial transcriptomic clusters (Fig. 1), with spatial “fibroblast-enriched signatures” highlighted.

**Extended Data Fig. 4:**
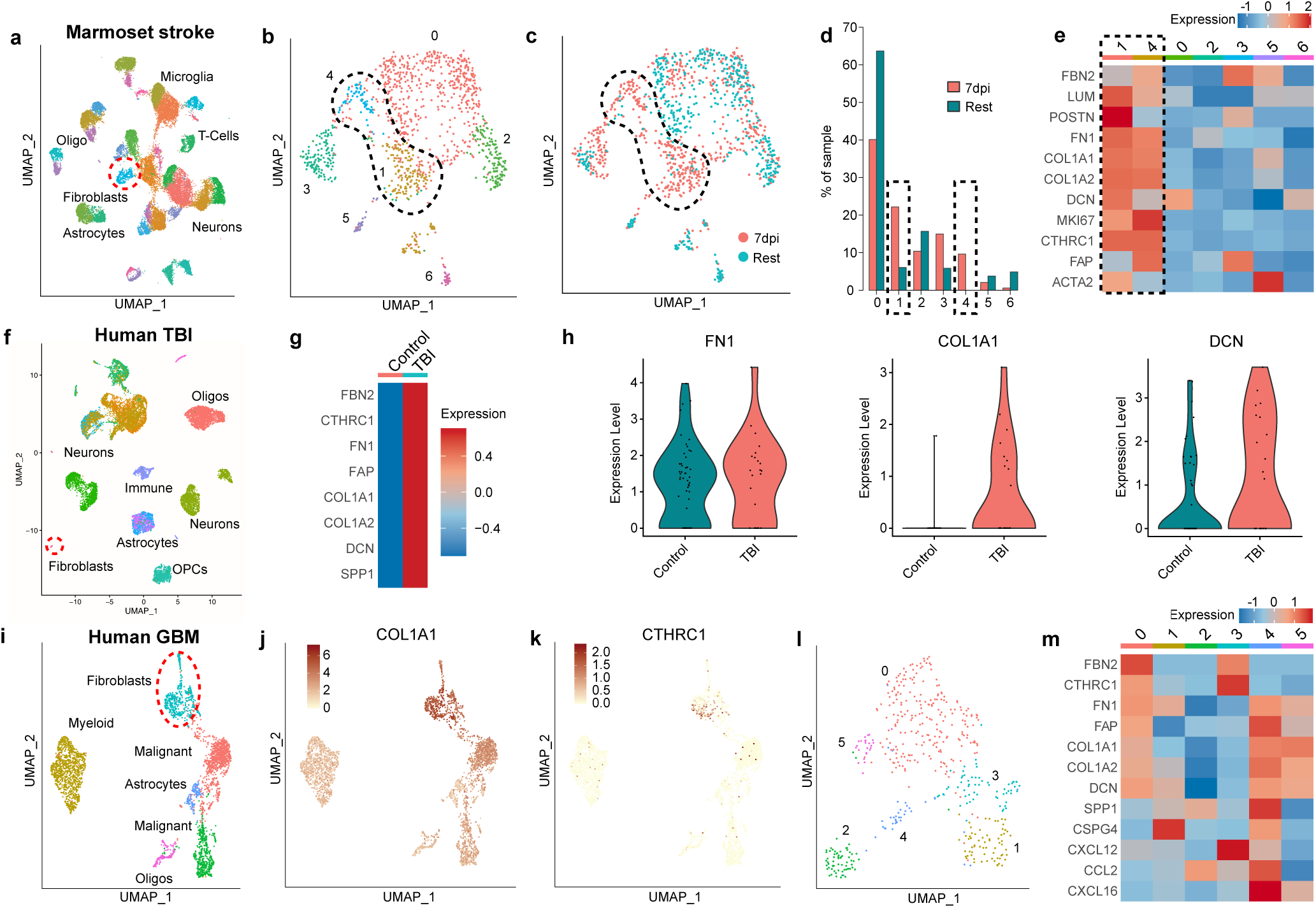
Transcriptomic identification of myofibroblast responses across species and injury models, related to Fig. 2. **a** Adult marmoset annotated snRNAseq data (Boghdadi et al.^76^) from endothelin-1-induced stroke (cortex, 7dpi). Fibroblasts highlighted (red dotted line) based on COL1A2 expression. **b-c**, UMAPs showing integrated fibroblasts from 7dpi (described in **a**) and resting adult marmoset dataset, showing annotated fibroblast subclusters (**b**) and timepoint (**c**). Black dotted line highlights 7dpi-enriched clusters 1 and 4. **d-e**, Bar plot showing relative cluster abundance (**d**, as proportion of fibroblast nuclei) and heatmap of select fibroblast/myofibroblast-related genes (**e**) across fibroblast clusters. Black dotted line highlights 7dpi-enriched clusters 1 and 4. **f**, Human TBI annotated snRNAseq data (Garza et al.^94^), including control tissues (varying timepoints). Fibroblasts highlighted (red dotted line) based on COL1A2 expression. **g-h**, Heatmap (**g**) and violin plots (**h**) showing elevated expression of selected fibroblast/myofibroblast genes in post-TBI fibroblasts. **i-k**, Human glioblastoma multiforme (GBM) tissue snRNAseq data (Jain et al.^16^), shown as UMAP (**i**) and feature plots of COL1A1 (**j**) or CTHRC1 (**k**). Fibroblasts are highlighted in (**i**) (red dotted line). **l-m**, UMAP showing reclustered fibroblasts (**l**) and heatmap showing fibroblast expression of select fibroblast/myofibroblast genes (**m**), enriched in cluster 0.

**Extended Data Fig. 5:**
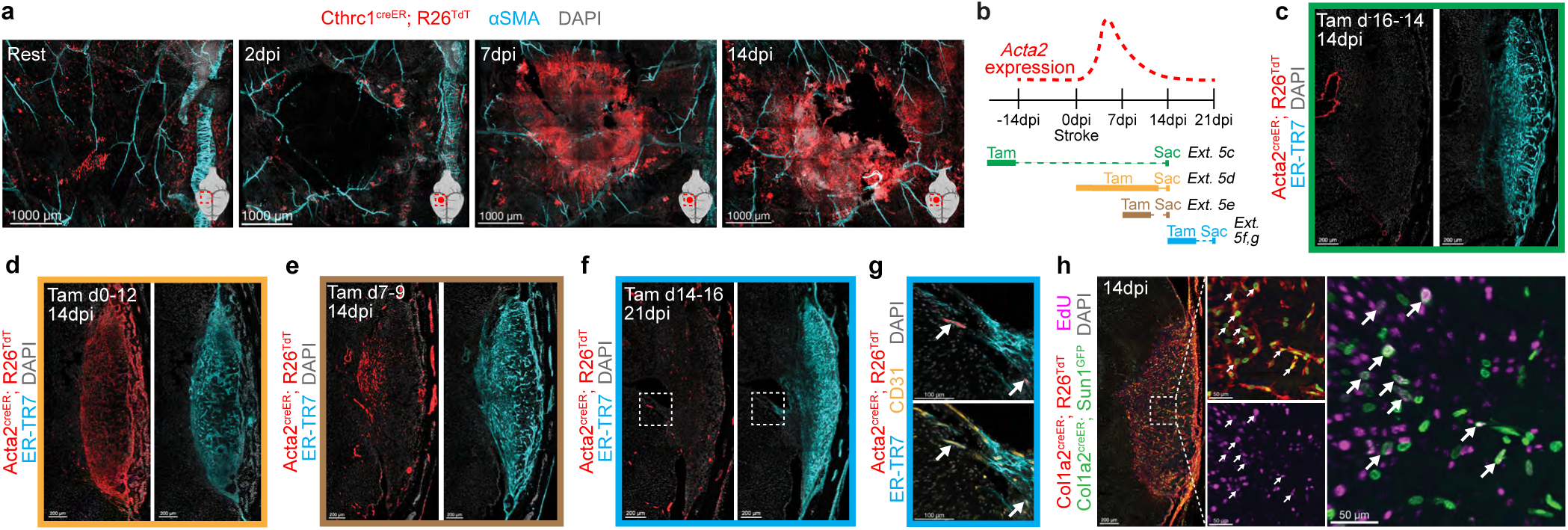
Additional *in vivo* validation of early TGFb-associated myofibroblasts, related to Fig. 2. **a** Confocal microscopy showing Cthrc1^creER^; Rosa26^TdT+^ fibroblasts in wholemounted dural meninges at indicated timepoints. **b**, Schematic showing tamoxifen induction for (**c-g**) and hypothesized trajectory for *Acta2* (gene for aSMA) expression. **c-g**, Confocal microscopy showing *Acta2-*lineage^+^ lesional fibroblasts after PT injury (tamoxifen induction and harvest days indicated), with lack of lineage-traced fibroblasts (**c**) but robust injury-induced *Acta2* (*TdT*) expression (**d**). Active *Acta2* expression is reduced but present at 7dpi (**e**, shown at 14dpi) and absent by 14dpi (**f**, shown at 21dpi). **g** (inset from **f**) shows expected Acta2^creER^; Rosa26^TdT+^ vascular smooth muscle cells (white arrows) near CD31^+^ vessels. 14μm slices; images represent 2 or more mice.

**Extended Data Fig. 6:**
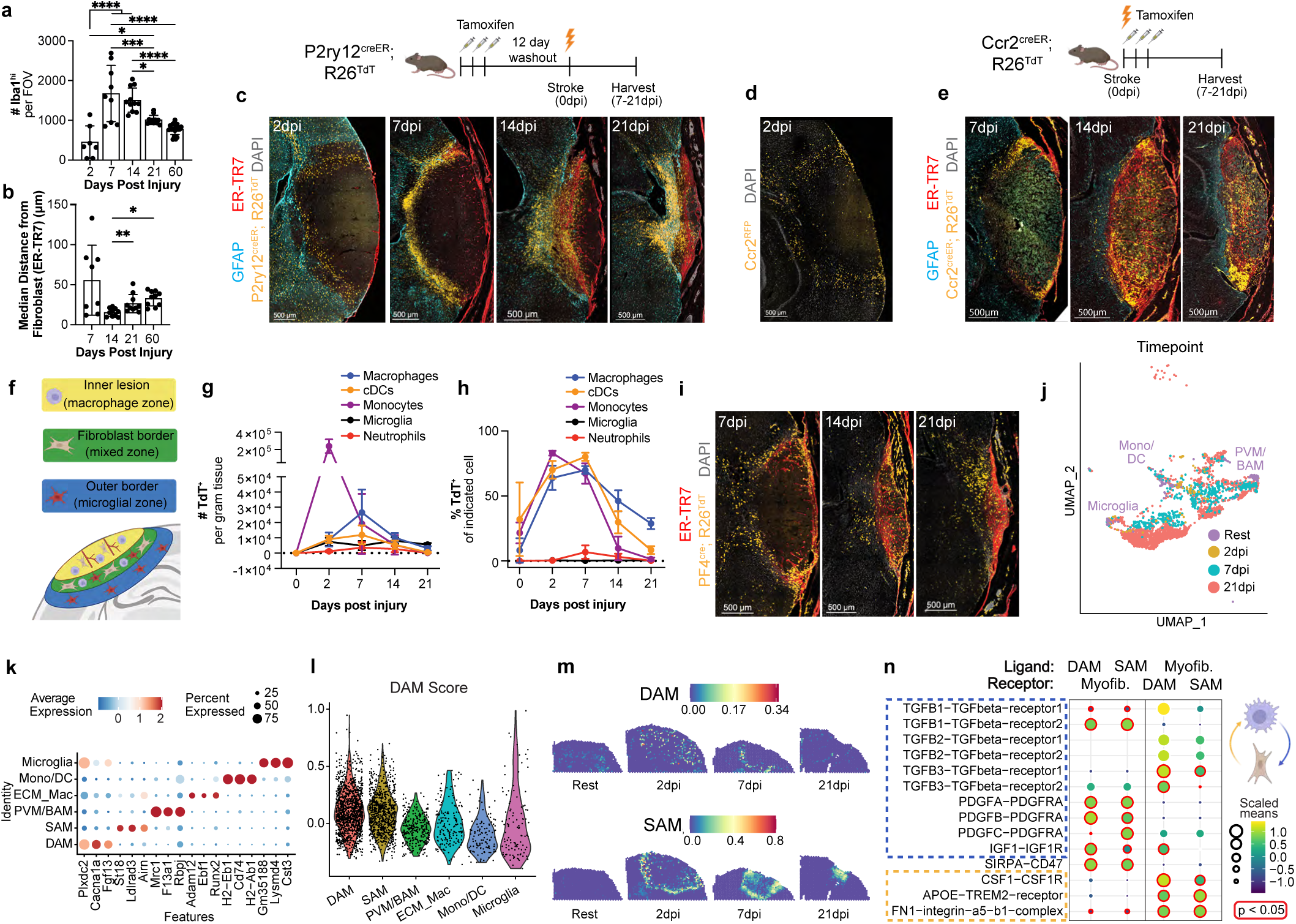
Additional characterization of injury-responsive profibrotic CNS myeloid cells, related to Fig. 2. **a-b** Quantification of Fig. 2o showing number of myeloid cells (Iba1^hi^, macrophages and reactive microglia) per slice (**a**) and median myeloid cell distance from the nearest fibroblast-associated ECM (ER-TR7^+^) surface (**b**). In **a**, n=7 slices from 3 mice (2dpi), 9 slices from 4 mice (7dpi), 11 slices from 5 mice (14/21dpi), or 20 slices from 5 mice (60dpi). In **b**, n=4 (7dpi) or 10 (14/21/60dpi) slices (2 slices/mouse). **c-f**, Immunofluorescent imaging from microglia lineage tracer mice (**c,** P2ry12^creER^; Rosa26^TdT^; tamoxifen days ^-^14-^-^12 before injury), or monocyte reporter/lineage tracer mice (**d**, Ccr2^RFP^; **e**, Ccr2^creER^; Rosa26^TdT^; tamoxifen 0-2dpi). Schematic (**f**) summarizes myeloid distribution by ontogeny. **g-h**, Time course showing *Ccr2-*lineage traced (Ccr2^TdT+^) cells as normalized counts (**g**) or as a proportion of indicated population (**h**). Tamoxifen 0-2dpi. n=4 (0/2dpi), 3 (7/21dpi), or 5 (14dpi) mice. **i**, Immunofluorescent imaging from BAM/PVM lineage tracer mice (PF4^cre^; Rosa26^TdT^). *PF4*-lineage^+^ cells (yellow) shown in perilesional regions (7dpi) or perilesional, core, and border regions (14/21dpi). **j-m**, UMAP of reclustered myeloid cells (**j**, colored by timepoint; major resting populations labelled), dot plot showing myeloid subcluster marker genes (**k**), violin plot of Disease Associated Microglia (“DAM”) score (**l**, an aggregate of 30 published DAM-specific markers), and DAM/SAM snRNAseq identities mapped onto spatial transcriptomic data (**m**). **n**, Ligand-receptor interactions between SAM or DAM and myofibroblasts (CellPhoneDB). Left half: ligand (first molecule in labelled pair) expressed by SAM/DAM, receptor expressed by myofibroblast. Right half: ligand expressed by myofibroblast, receptor expressed by SAM/DAM. Dotted lines and schematic (right) highlight modules with macrophage-expressed ligands (blue) or myofibroblast-expressed ligands (orange). Graphs show mean±s.d. ns=not significant, *p<0.05, **p<0.01, ***p<0.001, ****p<0.0001; one-way ANOVA, Tukey post-test. 14μm slices; images represent 2 or more mice.

**Extended Data Fig. 7:**
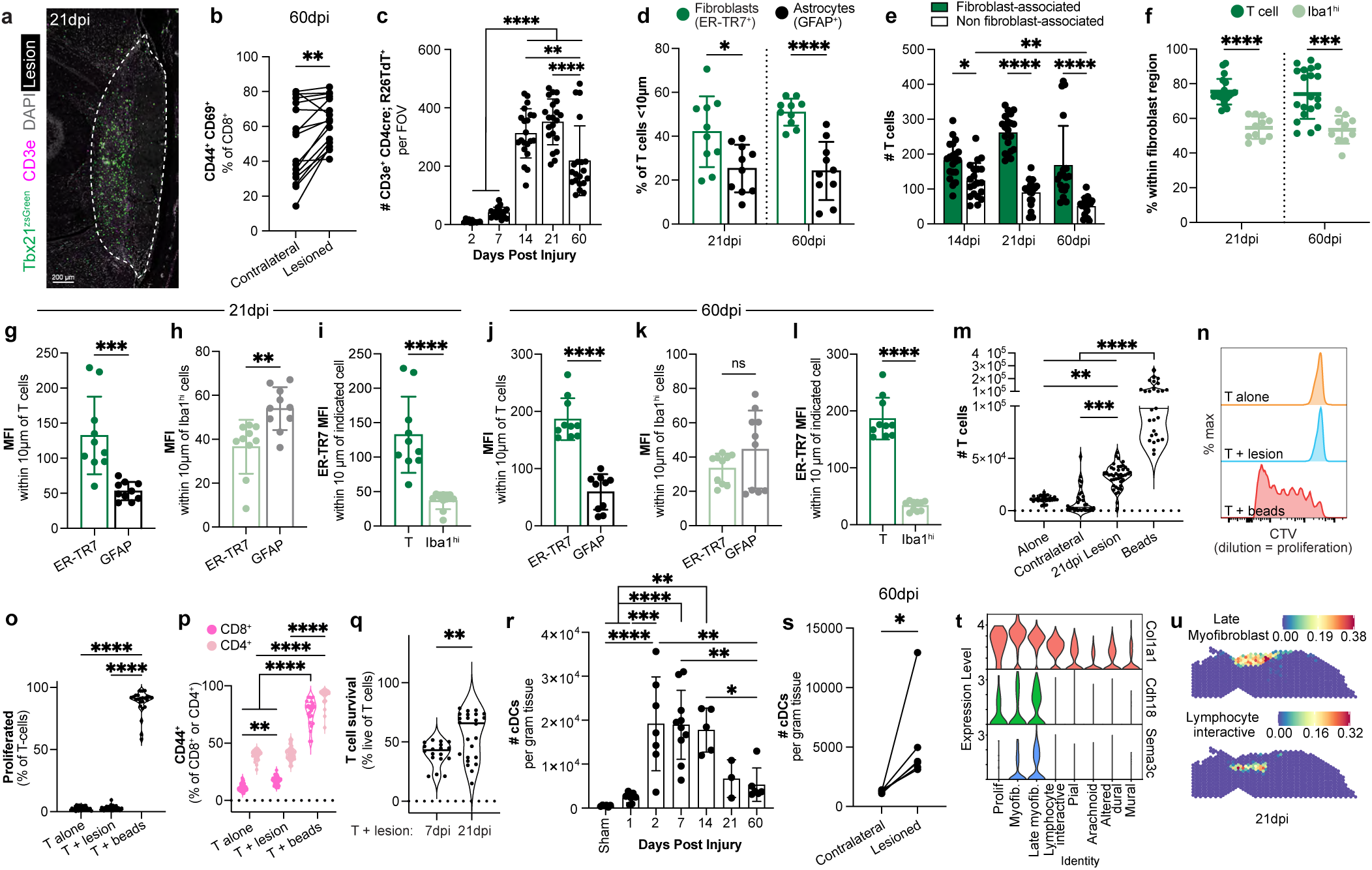
Additional characterization of fibroblast-lymphocyte interactions, related to Fig. 3. **a** Lesional type 1 lymphocytes (Tbet^zsGreen+^, 21dpi). **b**, Tissue resident memory CD8^+^ T cell distribution (CD44^+^ CD69^+^; 60dpi). n=17 mice. **c**, Perilesional T cell time course (CD4^cre^; Rosa26^TdT+^; CD3e^+^). n=11 slices from 3 mice (2dpi), 17 slices from 4 mice (7dpi), or 21 (14/21dpi) or 20 (60dpi) slices from 5 mice. **d-f**, T cell proportion <10μm from nearest fibroblast-associated ECM (ER-TR7^+^) or astrocyte (GFAP^+^) surface (**d**), T cells within or outside fibroblast-dense lesion (**e**), and T or myeloid cell (Iba1^hi^) proportion within lesion (**f**). n=10 slices (**d**, 2 slices/mouse); 20 slices (**e**, 4 slices/mouse); or 20 (T), 11 (Iba1^hi^/21dpi), or 10 (Iba1^hi^/60dpi) slices from 5 mice (**f**). **g-l**, Mean fluorescence intensity (MFI) of ER-TR7 (fibroblasts) and GFAP (astrocytes) within 10μm of each T (**g,j**) or myeloid cell (**h,k**), or ER-TR7 MFI comparison (**i,l**). 21dpi (**g-i**) or 60dpi (**j-l**). n=10 slices/timepoint (2 slices/mouse). **m-p**, T cell counts (**m**), proliferation (CTV dilution, **n**,**o**), and activation (**p**, CD44^+^) after coculture (21dpi lesions). n=30 (alone), 31 (contralateral), 44 (21dpi lesion), or 27 (beads) wells (**m**); or 27 (alone), 44 (lesion), or 24 (beads) wells (**o**,**p**). **q**, T cell survival after coculture with 7dpi or 21dpi lesion. n=18 (7dpi) or 24 (21dpi) wells. **r-s**, Conventional dendritic cell (cDC) cortical infiltration (**r**) and persistence (**s**) after PT injury. n=6 (sham/60dpi), 7 (1/2dpi), 10 (7dpi), 5 (14dpi), or 3 (21dpi) mice. **t**, Violin plots showing ECM-and smooth muscle-related genes^96,97^ (enriched among late myofibroblasts). **u**, Spatial plots showing late myofibroblast or lymphocyte interactive fibroblast distribution. Graphs show mean±s.d. ns=not significant, *p<0.05, **p<0.01, ***p<0.001, ****p<0.0001; paired Student’s T-test (**b**,**g-l,s**); oneway ANOVA, Tukey post-test (**c**,**m**,**o**,**r**); multiple T-tests, Holm-Sidak correction (**d**,**f**); two-way ANOVA, Sidak’s post-test (**e,p**); student’s T-test (**q**). 14μm slices; images represent two or more mice.

**Extended Data Fig. 8:**
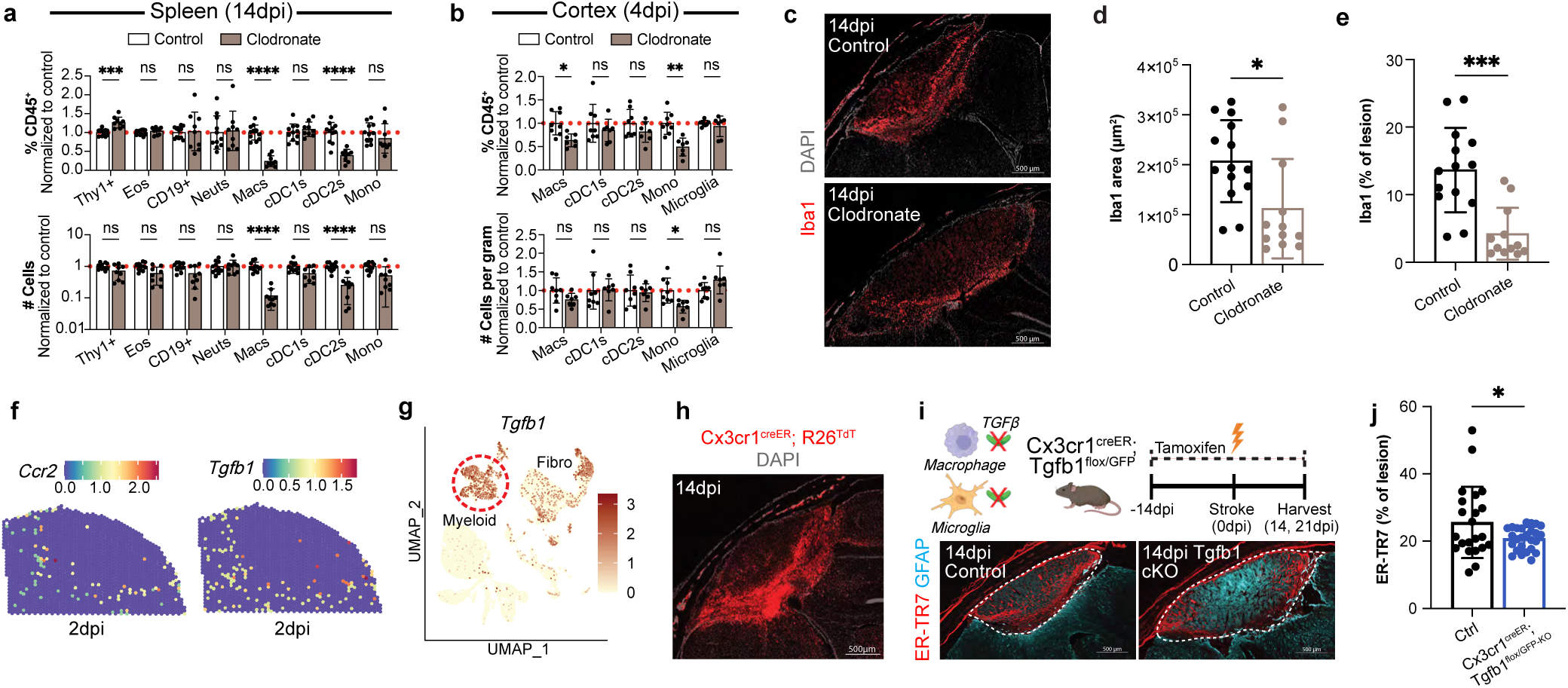
Profibrotic macrophages promote CNS fibroblast expansion, related to Fig. 4. **a-b** 14dpi splenic immune cells (**a**) or 4dpi cortical phagocytic myeloid cells (**b**) after treatment with control or clodronate liposomes, as frequency (top) or counts (bottom). n=11 (control) or 9 (clodronate) 11 mice (**a**) or 8 (control) or 7 (clodronate) mice (**b**). Within each experiment, clodronate liposome-treated mice were normalized to control liposome-treated mice. **c-e**, Perilesional myeloid cell (Iba1) staining at 14dpi, with images (**c**) and quantification via thresholding (**d**) and normalization to reactive astrocyte (GFAP)-traced lesion area (**e**). n=14 (control) or 12 (clodronate) slices (2 slices/mouse). **f**, Spatial transcriptomic feature plots from Visium dataset (Fig. 1) showing *Ccr2* and *Tgfb1* transcripts in perilesional distribution at 2dpi. **g**, Feature plot of *Tgfb1* from mouse snRNAseq (plotted in order of expression); dotted line highlights myeloid cells. **h**, Representative image showing recombined macrophages and microglia (TdT^+^) in a Cx3cr1^creER^; Rosa26^TdT^ mouse (tamoxifen as below; 14dpi). **i-j**, Lesional images (**i**; dotted line shows lesion boundary) and ER-TR7 coverage (**j**) following genetic targeting of macrophage and microglial *Tgfb1* in Cx3cr1^creER^; Tgfb1^flox/GFP-KO^ mice (14dpi). Controls were littermate Cx3cr1^creER^; Tgfb1^flox/+^ mice; tamoxifen as in Fig. 4d. n=22 (control) or 26 (*Tgfb1* cKO) slices (2 slices/mouse). Graphs show mean±s.d. ns=not significant, *p<0.05, **p<0.01, ***p<0.001, ****p<0.0001; multiple T-tests, Holm-Sidak correction (**a**,**b**); Student’s T-test (**d**,**e**,**j**). 14μm slices; images represent two or more mice.

**Extended Data Fig. 9:**
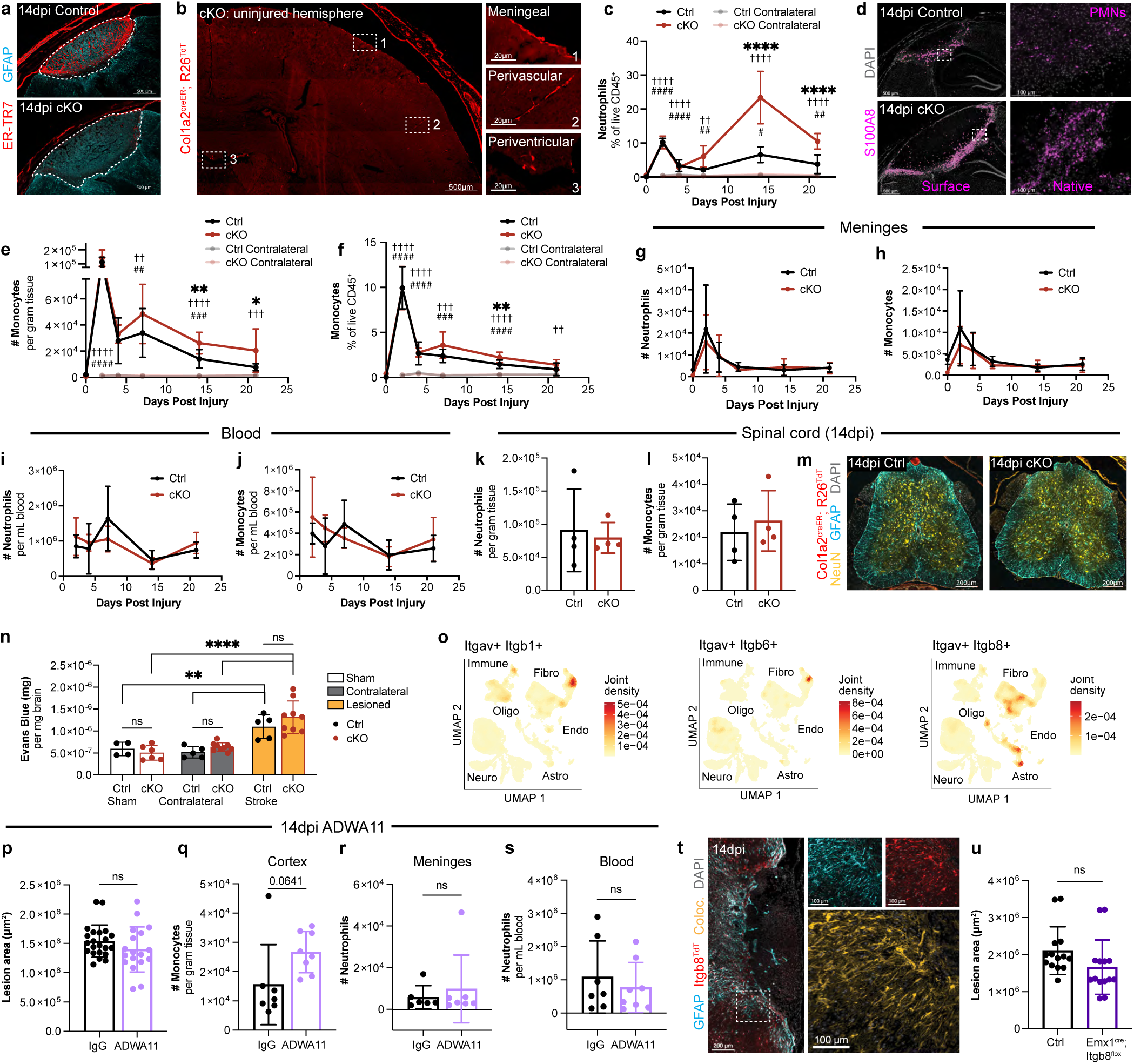
TGFb drives CNS wound healing and resolution of inflammation, related to Fig. 4. **a** Lesional images showing ER-TR7 in controls or cKOs (Col1a2^creER^; Tgfbr2^flox^), 14dpi (dotted line shows lesion boundary). **b**, Intact fibroblast (Col1a2^creER^; Rosa26^TdT+^) topography in uninjured cKO parenchyma (insets: leptomeningeal/perivascular/periventricular fibroblasts). **c**, Time course showing cortical neutrophils in controls/cKOs. n=6 (0dpi-control [unrelated], 2/4/7dpi-control, 2/14/21dpi-cKO), 8 (14dpi-control), 9 (21dpi-control), 3 (0dpi-cKO), or 5 (4dpi-cKO) mice. **d**, S100A8^+^ neutrophils (PMN; left, surfaced; right, native) in controls (top) or cKOs (bottom), 14dpi. **e-f**, Time course showing cortical monocyte frequency (**e**) or normalized counts (**f**) in controls/cKOs. n as in (**c**). **g-l**, Neutrophil or monocyte counts in meninges (**g-h**), blood (**i-j**), or 14dpi spinal cord (**k**-**l**) of controls/cKOs. n=3 (0dpi control [unrelated]/cKO), 6 (2/4/7dpi control, 2/14/21dpi cKO), 8 (14dpi control), 9 (21dpi control), 5 (7dpi cKO), and 5 (4dpi cKO, **g-h**) or 6 (4dpi cKO, **i-j**) mice; n=4 mice/group (**k**-**l**). **m**, Control/cKO spinal cords showing intact neurons without fibrosis/gliosis (14dpi). **n**, Evans Blue extravasation in sham or contralateral/lesioned hemispheres of controls/cKOs, 4dpi. n=4 (shamcontrol), 6 (sham-cKO), 5 (contralateral/lesioned-control), or 9 (contralateral/lesioned-cKO) mice. **o**, Co-expression of TGFb-activating integrin pairs *Itgav* and *Itgb1, Itgb6,* or *Itgb8* (snRNAseq). **p-s**, Lesion size (**p**), cortical monocytes (**q**), and meningeal (**r**) or blood neutrophils (**s**) in IgG-and ADWA11-treated mice (14dpi). n=22 (IgG) or 18 (ADWA11) slices (**p**, 2 slices/mouse); n=7 (IgG) or 8 (ADWA11) mice (**q**,**s**) or 6 (IgG) or 7 (ADWA11) mice (**r**). **t**, Itgb8^TdT^ mouse, with perilesional/astrocytic *TdT*, 14dpi (insets: GFAP/TdT colocalization, yellow). **u**, Lesion size (14dpi) in Emx1^cre^; Itgb8^flox^ mice and controls. n=14 slices/group (2 slices/mouse). Graphs show mean±s.d. ns=not significant, *p<0.05, **p<0.01, ***p<0.001, ****p<0.0001; * lesioned-control vs. lesioned-cKO; # lesioned-control vs. contralateralcontrol; ^†^ lesioned-cKO vs. contralateral-cKO (**c**,**e**,**f**). Two-way ANOVA, Sidak’s post-test (**c**,**e-f** [per injury timepoint], **g-j**,**n**). Student’s T-test (**k**,**l**,**p-s**,**u**). 14μm slices; images represent two or more mice.

**Extended Data Fig. 10:**
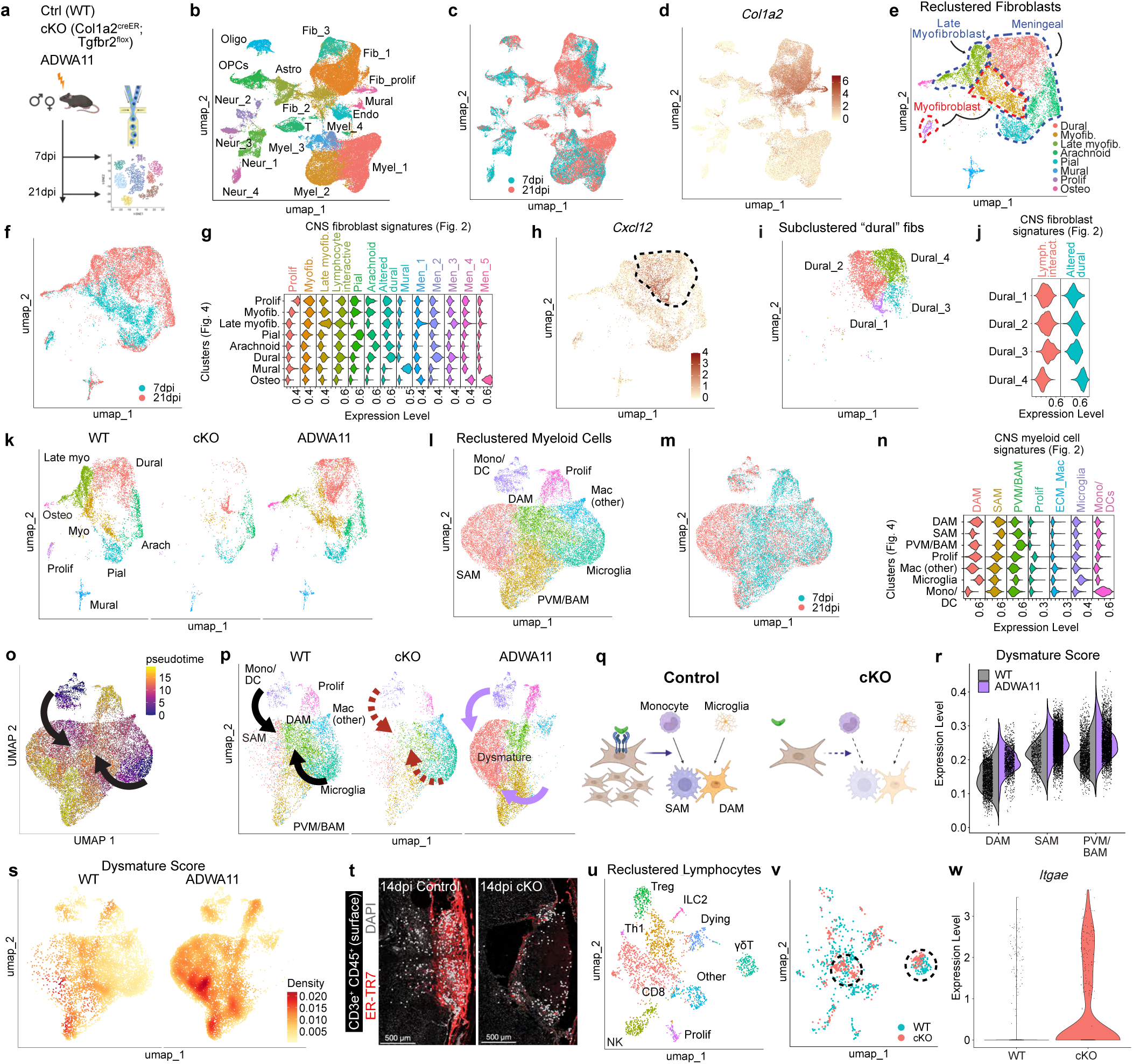
Additional molecular characterization of injury response after fibroblast impairment, related to Fig. 4. **a** snRNAseq schematic. WT, cKO (Col1a2^creER^; Tgfbr2^flox^), and a^v^b^8^-blocking antibody (ADWA11)-treated mice were harvested at 7 and 21dpi (2 mice per timepoint/condition), lesions were micro-dissected, and nuclei were sorted (DAPI^+^) and sequenced. **bd**, Global UMAPs from snRNAseq data showing indicated cell types (**b**), timepoint metadata (**c**), or *Col1a1* expression (**d**). **e-f**, UMAPs showing fibroblast subclusters (**e**) or timepoint (**f**). **g**, Violin plots mapping “CNS fibroblast signatures” (derived from Fig. 2, x-axis) onto clusters identified in **e** (y-axis). **h-j**, *Cxcl12* expression among fibroblasts (**h**, “dural” fibroblasts highlighted), UMAP showing subclustered “dural” fibroblasts (**i**), and violin plots mapping expression of “Lymphocyte interactive” and “Altered dural” signatures (derived from Fig. 2) onto “dural fibroblast” subclusters (**j**), highlighting 3 lymphocyte-interactive-like and 1 altered-dural-like subcluster(s). **k**, Fibroblast UMAP separated by condition, with pan-fibroblast reduction in cKO mice and late myofibroblast reduction in a^v^b^8^-blocked (ADWA11) mice. **l-m**, UMAPs showing myeloid cell subclusters (**l**) or timepoint (**m**). **n**, Violin plots mapping expression of “CNS myeloid cell signatures” (derived from Fig. 2, x-axis) onto clusters identified in **l** (y-axis). **o-p**, Pseudotime (**o**, Monocle3) and UMAP plots separated by condition (**p**) showing potential myeloid differentiation trajectories, including monocyte-to-SAM (top) and microgliato-DAM (bottom), with monocytes and microglia as root states. **q**, Schematic showing effect of TGFbdriven fibroblasts on SAM/DAM formation. **r-s**, Violin (**r**) or UMAP plots (**s**) showing “dysmature” transcriptional signature across conditions. **t**, Surfaced T cells (CD3e^+^ CD45^+^) in control and cKO mice, with fibroblast-rich (ER-TR7^+^) lesion. **u-v**, UMAP showing lymphocyte subclusters (**u**) or genotype (**v**, WT and cKO). Dotted black lines highlight CD8 and gdT cells. **w**, Violin plot of lymphocyte *Itgae* expression.

## References

1. Iadecola, C., Buckwalter, M. S. & Anrather, J. Immune responses to stroke: mechanisms, modulation, and therapeutic potential. J Clin Invest 130, 2777–2788 (2020).

2. Fehily, B. & Fitzgerald, M. Repeated Mild Traumatic Brain Injury. Cell Transplant 26, 1131–1155 (2017).

3. Karsy, M. & Hawryluk, G. Modern Medical Management of Spinal Cord Injury. Curr Neurol Neurosci Rep 19, 65 (2019).

4. Al-Qazzaz, N. K., Ali, S. H., Ahmad, S. A., Islam, S. & Mohamad, K. Cognitive impairment and memory dysfunction after a stroke diagnosis: a post-stroke memory assessment. Neuropsychiatr Dis Treat 10, 1677–1691 (2014).

5. Stephanie S. Holden et al. Complement factor C1q mediates sleep spindle loss and epileptic spikes after mild brain injury. Science 373, eabj2685 (2021).

6. Brett, B. L., Gardner, R. C., Godbout, J., Dams-O’Connor, K. & Keene, C. D. Traumatic Brain Injury and Risk of Neurodegenerative Disorder. Biol Psychiatry 91, 498–507 (2022).

7. Emberson, J. et al. Effect of treatment delay, age, and stroke severity on the effects of intravenous thrombolysis with alteplase for acute ischaemic stroke: a meta-analysis of individual patient data from randomised trials. The Lancet 384, 1929– 1935 (2014).

8. Alawieh, A., Zhao, J. & Feng, W. Factors affecting post-stroke motor recovery: Implications on neurotherapy after brain injury. Behav Brain Res 340, 94–101 (2018).

9. Zehendner, C. M. et al. Traumatic brain injury results in rapid pericyte loss followed by reactive pericytosis in the cerebral cortex. Scientific Reports 5, 13497 (2015).

10. Dias, D. O. et al. Reducing Pericyte-Derived Scarring Promotes Recovery after Spinal Cord Injury. Cell 173, 153–165.e22 (2018).

11. Dias, D. O. et al. Pericyte-derived fibrotic scarring is conserved across diverse central nervous system lesions. Nature Communications 12, 5501 (2021).

12. Xu, L., et al. Fibroblasts repair blood-brain barrier damage and hemorrhagic brain injury via TIMP2. Cell Reports 41, (2022).

13. Fernández-Klett, F. et al. Early loss of pericytes and perivascular stromal cell-induced scar formation after stroke. J Cereb Blood Flow Metab 33, 428–439 (2013).

14. Dorrier, C. E., Jones, H. E., Pintarić, L., Siegenthaler, J. A. & Daneman, R. Emerging roles for CNS fibroblasts in health, injury and disease. Nature Reviews Neuroscience (2021) doi:10.1038/s41583-021-00525-w.

15. Pikor, N. B. et al. Integration of Th17-and Lymphotoxin-Derived Signals Initiates Meningeal-Resident Stromal Cell Remodeling to Propagate Neuroinflammation. Immunity 43, 1160–73 (2015).

16. Jain, S. et al. Single-cell RNA sequencing and spatial transcriptomics reveal cancer-associated fibroblasts in glioblastoma with protumoral effects. J Clin Invest 133, e147087.

17. Månberg, A. et al. Altered perivascular fibroblast activity precedes ALS disease onset. Nature Medicine 27, 640–646 (2021).

18. Dorrier, C. E. et al. CNS fibroblasts form a fibrotic scar in response to immune cell infiltration. Nature Neuroscience 24, 234–244 (2021).

19. Buechler, M. B. et al. Cross-tissue organization of the fibroblast lineage. Nature (2021) doi:10.1038/s41586-021-03549-5.

20. Kinchen, J. et al. Structural Remodeling of the Human Colonic Mesenchyme in Inflammatory Bowel Disease. Cell 175, 372–386.e17 (2018).

21. Boothby, I. C. et al. Early-life inflammation primes a T helper 2 cell–fibroblast niche in skin. Nature (2021) doi:10.1038/s41586-021-04044-7.

22. Xu, Z. et al. Anatomically distinct fibroblast subsets determine skin autoimmune patterns. Nature (2021) doi:10.1038/s41586-021-04221-8.

23. Dahlgren, M. W. et al. Adventitial Stromal Cells Define Group 2 Innate Lymphoid Cell Tissue Niches. Immunity 50, 707–722.e6 (2019).

24. Hu, K. H. et al. Transcriptional space-time mapping identifies concerted immune and stromal cell patterns and gene programs in wound healing and cancer. Cell Stem Cell 30, 885–903.e10 (2023).

25. Tsukui, T. et al. Collagen-producing lung cell atlas identifies multiple subsets with distinct localization and relevance to fibrosis. Nature Communications 11, 1920– 1920 (2020).

26. Fabre, T. et al. Identification of a broadly fibrogenic macrophage subset induced by type 3 inflammation. Science Immunology 8, eadd8945 (2023).

27. Rustenhoven, J. et al. Functional characterization of the dural sinuses as a neuroimmune interface. Cell (2021) 10.1016/j.cell.2020.12.040.

28. Mastorakos, P. & McGavern, D. The anatomy and immunology of vasculature in the central nervous system. Science Immunology 4, eaav0492 (2019).

29. Frangogiannis, N. Transforming growth factor-β in tissue fibrosis. The Journal of experimental medicine 217, e20190103–e20190103 (2020).

30. Tsukui, T. & Sheppard, D. Tracing the origin of pathologic pulmonary fibroblasts. bioRxiv 2022.11.18.517147 (2022) doi:10.1101/2022.11.18.517147.

31. Hoeft, K., et al. Platelet-instructed SPP1+ macrophages drive myofibroblast activation in fibrosis in a CXCL4-dependent manner. Cell Reports 42, (2023).

32. Drieu, A. et al. Parenchymal border macrophages regulate the flow dynamics of the cerebrospinal fluid. Nature (2022) doi:10.1038/s41586-022-05397-3.

33. Keren-Shaul, H. et al. A Unique Microglia Type Associated with Restricting Development of Alzheimer’s Disease. Cell 169, 1276–1290.e17 (2017).

34. Mastorakos, P. et al. Temporally distinct myeloid cell responses mediate damage and repair after cerebrovascular injury. Nature Neuroscience 24, 245–258 (2021).

35. Griffith, J. W., Sokol, C. L. & Luster, A. D. Chemokines and chemokine receptors: positioning cells for host defense and immunity. Annu Rev Immunol 32, 659–702 (2014).

36. DeSisto, J. et al. Single-Cell Transcriptomic Analyses of the Developing Meninges Reveal Meningeal Fibroblast Diversity and Function. Developmental Cell 54, 43–59.e4 (2020).

37. Derynck, R. & Budi, E. H. Specificity, versatility, and control of TGF-β family signaling. Science Signaling 12, eaav5183 (2019).

38. Henderson, N. C. et al. Targeting of αv integrin identifies a core molecular pathway that regulates fibrosis in several organs. Nat Med 19, 1617–1624 (2013).

39. Yin, Z. et al. APOE4 impairs the microglial response in Alzheimer’s disease by inducing TGFβ-mediated checkpoints. Nat Immunol 24, 1839–1853 (2023).

40. Arnold, T. D. et al. Impaired αVβ8 and TGFβ signaling lead to microglial dysmaturation and neuromotor dysfunction. Journal of Experimental Medicine 216, 900–915 (2019).

41. McKinsey, G. L. et al. Radial glia control microglial differentiation via integrin avb8-dependent trans-activation of TGFB1. 2023.07.13.548459 Preprint at 10.1101/2023.07.13.548459 (2023).

42. Linda M. Wakim, Amanda Woodward-Davis & Michael J. Bevan. Memory T cells persisting within the brain after local infection show functional adaptations to their tissue of residence. Proceedings of the National Academy of Sciences 107, 17872– 17879 (2010).

43. Sommer, C. J. Ischemic stroke: experimental models and reality. Acta Neuropathol 133, 245–261 (2017).

44. Soderblom, C. et al. Perivascular fibroblasts form the fibrotic scar after contusive spinal cord injury. J Neurosci 33, 13882–13887 (2013).

45. Yang, A. C. et al. A human brain vascular atlas reveals diverse cell mediators of Alzheimer’s disease risk. bioRxiv 2021.04.26.441262 (2021) doi:10.1101/2021.04.26.441262.

46. Vanlandewijck, M. et al. A molecular atlas of cell types and zonation in the brain vasculature. Nature 554, 475–480 (2018).

47. Rustenhoven, J. & Kipnis, J. Brain borders at the central stage of neuroimmunology. Nature 612, 417–429 (2022).

48. Krishnamurty, A. T. et al. LRRC15+ myofibroblasts dictate the stromal setpoint to suppress tumour immunity. Nature 611, 148–154 (2022).

49. Öhlund, D. et al. Distinct populations of inflammatory fibroblasts and myofibroblasts in pancreatic cancer. The Journal of experimental medicine 214, 579–596 (2017).

50. Salmon, H. et al. Matrix architecture defines the preferential localization and migration of T cells into the stroma of human lung tumors. J Clin Invest 122, 899– 910 (2012).

## Supplemental references

51. He, L. et al. Preexisting endothelial cells mediate cardiac neovascularization after injury. J Clin Invest 127, 2968–2981 (2017).

52. Krempen, K. et al. Far upstream regulatory elements enhance position-independent and uterus-specific expression of the murine alpha1(I) collagen promoter in transgenic mice. Gene Expr 8, 151–163 (1999).

53. Hamilton, T. G., Klinghoffer, R. A., Corrin, P. D. & Soriano, P. Evolutionary divergence of platelet-derived growth factor alpha receptor signaling mechanisms. Mol Cell Biol 23, 4013–4025 (2003).

54. Rivers, L. E. et al. PDGFRA/NG2 glia generate myelinating oligodendrocytes and piriform projection neurons in adult mice. Nat Neurosci 11, 10.1038/nn.2220 (2008).

55. Šošić, D., Richardson, J. A., Yu, K., Ornitz, D. M. & Olson, E. N. Twist regulates cytokine gene expression through a negative feedback loop that represses NF-kappaB activity. Cell 112, 169–180 (2003).

56. Wendling, O., Bornert, J.-M., Chambon, P. & Metzger, D. Efficient temporally-controlled targeted mutagenesis in smooth muscle cells of the adult mouse. genesis 47, 14–18 (2009).

57. Lee, P. P. et al. A critical role for Dnmt1 and DNA methylation in T cell development, function, and survival. Immunity 15, 763–774 (2001).

58. Zhu, J. et al. The transcription factor T-bet is induced by multiple pathways and prevents an endogenous Th2 cell program during Th1 cell responses. Immunity 37, 660–673 (2012).

59. Li, M. O., Wan, Y. Y. & Flavell, R. A. T Cell-Produced Transforming Growth Factor-β1 Controls T Cell Tolerance and Regulates Th1-and Th17-Cell Differentiation. Immunity 26, 579–591 (2007).

60. Chytil, A., Magnuson, M. A., Wright, C. V. E. & Moses, H. L. Conditional inactivation of the TGF-β type II receptor using Cre:Lox. genesis 32, 73–75 (2002).

61. Nakawesi, J. et al. αvβ8 integrin-expression by BATF3-dependent dendritic cells facilitates early IgA responses to Rotavirus. Mucosal Immunol 14, 53–67 (2021).

62. Proctor, J. M., Zang, K., Wang, D., Wang, R. & Reichardt, L. F. Vascular Development of the Brain Requires β8 Integrin Expression in the Neuroepithelium. J. Neurosci. 25, 9940–9948 (2005).

63. Gorski, J. A. et al. Cortical excitatory neurons and glia, but not GABAergic neurons, are produced in the Emx1-expressing lineage. J Neurosci 22, 6309–6314 (2002).

64. Sudwarts, A. et al. BIN1 is a key regulator of proinflammatory and neurodegeneration-related activation in microglia. Mol Neurodegener 17, 33 (2022).

65. Cohen, J. N. et al. Regulatory T cells in skin mediate immune privilege of the hair follicle stem cell niche. Science Immunology 9, eadh0152 (2024).

66. Labat-gest, V. & Tomasi, S. Photothrombotic ischemia: a minimally invasive and reproducible photochemical cortical lesion model for mouse stroke studies. J Vis Exp 50370 (2013) doi:10.3791/50370.

67. Lee, J. K. et al. Photochemically induced cerebral ischemia in a mouse model. Surg Neurol 67, 620–5; discussion 625 (2007).

68. Liu, N.-W. et al. Evolutional Characterization of Photochemically Induced Stroke in Rats: a Multimodality Imaging and Molecular Biological Study. Translational Stroke Research 8, 244–256 (2017).

69. Gonzalez, F. F. et al. Erythropoietin increases neurogenesis and oligodendrogliosis of subventricular zone precursor cells after neonatal stroke. Stroke 44, 753–758 (2013).

70. McKinsey, G. L. et al. A new genetic strategy for targeting microglia in development and disease. eLife 9, e54590 (2020).

71. Larpthaveesarp, A. & Gonzalez, F. F. Transient Middle Cerebral Artery Occlusion Model of Neonatal Stroke in P10 Rats. J Vis Exp 54830 (2017) doi:10.3791/54830.

72. Teo, L. et al. Replicating infant-specific reactive astrocyte functions in the injured adult brain. Prog Neurobiol 204, 102108 (2021).

73. Teo, L. & Bourne, J. A. A Reproducible and Translatable Model of Focal Ischemia in the Visual Cortex of Infant and Adult Marmoset Monkeys. Brain Pathology 24, 459– 474 (2014).

74. Stockis, J. et al. Blocking immunosuppression by human Tregs in vivo with antibodies targeting integrin αVβ8. Proc Natl Acad Sci U S A 114, E10161–E10168 (2017).

75. Dodagatta-Marri, E. et al. Integrin αvβ8 on T cells suppresses anti-tumor immunity in multiple models and is a promising target for tumor immunotherapy. Cell Reports 36, 109309 (2021).

76. Boghdadi, A. G. et al. NogoA-expressing astrocytes limit peripheral macrophage infiltration after ischemic brain injury in primates. Nature Communications 12, 6906 (2021).

77. Renier, N. et al. iDISCO: A Simple, Rapid Method to Immunolabel Large Tissue Samples for Volume Imaging. Cell 159, 896–910 (2014).

78. Slyper, M. et al. A single-cell and single-nucleus RNA-Seq toolbox for fresh and frozen human tumors. Nat Med 26, 792–802 (2020).

79. Galatro, T. F., Vainchtein, I. D., Brouwer, N., Boddeke, E. W. G. M. & Eggen, B. J. L. Isolation of Microglia and Immune Infiltrates from Mouse and Primate Central Nervous System. Methods Mol Biol 1559, 333–342 (2017).

80. Nott, A., Schlachetzki, J. C. M., Fixsen, B. R. & Glass, C. K. Nuclei isolation of multiple brain cell types for omics interrogation. Nat Protoc 16, 1629–1646 (2021).

81. Radu, M. & Chernoff, J. An in vivo Assay to Test Blood Vessel Permeability. J Vis Exp 50062 (2013) doi:10.3791/50062.

82. Hao, Y. et al. Integrated analysis of multimodal single-cell data. Cell 184, 3573–3587.e29 (2021).

83. Blighe, K., et al. EnhancedVolcano: Publication-ready volcano plots with enhanced colouring and labeling. Bioconductor version: Release (3.18) 10.18129/B9.bioc.EnhancedVolcano (2023).

84. Sbierski-Kind, J. et al. >Group 2 innate lymphoid cells constrain type 3/17 lymphocytes in shared stromal niches to restrict liver fibrosis. 2023.04.26.537913 Preprint at 10.1101/2023.04.26.537913 (2023).

85. Lenburg, M. E., Sinha, A., Faller, D. V. & Denis, G. V. Tumor-specific and Proliferation-specific Gene Expression Typifies Murine Transgenic B Cell Lymphomagenesis. J Biol Chem 282, 4803 (2007).

86. Whitfield, M. L., George, L. K., Grant, G. D. & Perou, C. M. Common markers of proliferation. Nat Rev Cancer 6, 99–106 (2006).

87. Korsunsky, I., Nathan, A., Millard, N. & Raychaudhuri, S. Presto scales Wilcoxon and auROC analyses to millions of observations. 653253 Preprint at 10.1101/653253 (2019).

88. Alquicira-Hernandez, J. & Powell, J. E. Nebulosa recovers single-cell gene expression signals by kernel density estimation. Bioinformatics 37, 2485–2487 (2021).

89. Marsh, S. samuel-marsh/scCustomize: Version 2.0.1. doi:10.5281/zenodo.10161832.

90. Wu, T. et al. clusterProfiler 4.0: A universal enrichment tool for interpreting omics data. Innovation (Camb) 2, 100141 (2021).

91. Browaeys, R., Saelens, W. & Saeys, Y. NicheNet: modeling intercellular communication by linking ligands to target genes. Nat Methods 17, 159–162 (2020).

92. Cable, D. M. et al. Robust decomposition of cell type mixtures in spatial transcriptomics. Nat Biotechnol 40, 517–526 (2022).

93. Garcia-Alonso, L. et al. Single-cell roadmap of human gonadal development. Nature 607, 540–547 (2022).

94. Garza, R. et al. Single-cell transcriptomics of resected human traumatic brain injury tissues reveals acute activation of endogenous retroviruses in oligodendroglia. 2022.09.07.506982 Preprint at 10.1101/2022.09.07.506982 (2023).

95. Korsunsky, I. et al. Fast, sensitive and accurate integration of single-cell data with Harmony. Nat Methods 16, 1289–1296 (2019).

96. Junghof, J. et al. CDH18 is a fetal epicardial biomarker regulating differentiation towards vascular smooth muscle cells. npj Regen Med 7, 1–14 (2022).

97. Lepore, J. J. et al. GATA-6 regulates semaphorin 3C and is required in cardiac neural crest for cardiovascular morphogenesis. J Clin Invest 116, 929–939 (2006).

