## Supplementary Information for "Dynamic fibroblast-immune interactions shape wound healing after brain injury"

**Supplementary Video 1:** Whole lesion (PT injury, 14dpi, iDISCO) with fibroblasts and associated ECM.

**Supplementary Video 2:** Thick section image (200µm, iDISCO; PT injury, 7dpi) showing vascular remodeling, with PDGFRa$^{GFP+}$ nuclei, marking both fibroblasts and oligodendrocyte precursor cells (OPCs), enriched in perivascular regions.

**Supplementary Video 3:** Persistent *Cthrc1*-lineage$^{+}$ fibroblasts and surrounding gliosis at 21dpi (14µm; PT injury, tamoxifen 0-21dpi).

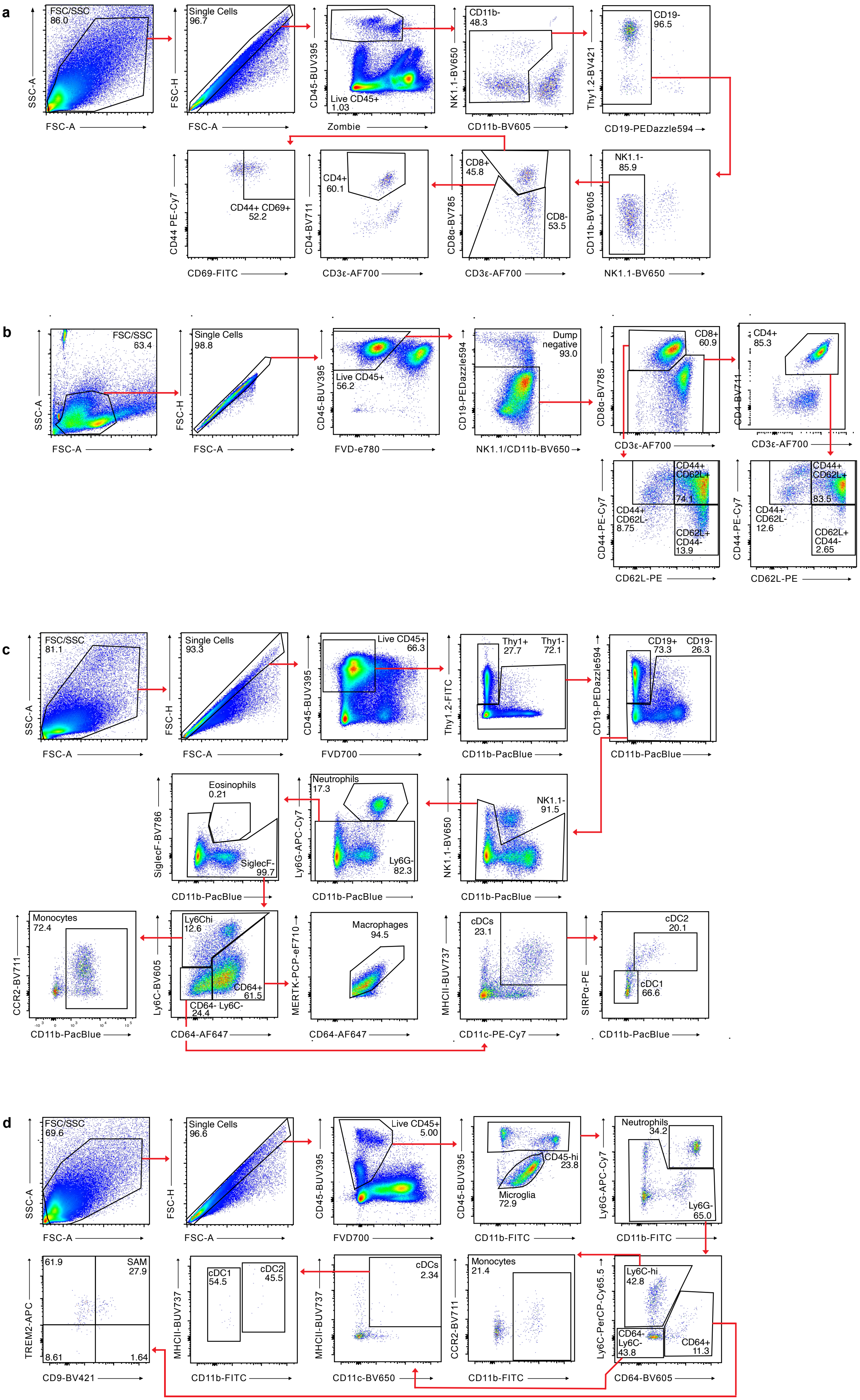
a
FSC/SSC 86.0
SSC-A
FSC-A
Single Cells 96.7
FSC-H
FSC-A
CD45-BUV395
Zombie
Live CD45+ 1.03
CD11b- 48.3
NK1.1-BV650
CD11b-BV605
CD19- 96.5
Thy1.2-BV421
CD19-PEDazzle594
CD44 PE-Cy7
CD69-FITC
CD44+ CD69+ 52.2
CD4+ 60.1
CD4-BV711
CD3ε-AF700
CD8+ 45.8
CD8α-BV785
CD3ε-AF700
CD8- 53.5
NK1.1- 85.9
CD11b-BV605
NK1.1-BV650
b
FSC/SSC 63.4
SSC-A
FSC-A
Single Cells 98.8
FSC-H
FSC-A
CD45-BUV395
FVD-e780
Live CD45+ 56.2
Dump negative 93.0
CD19-PEDazzle594
NK1.1/CD11b-BV650
CD8+ 60.9
CD8α-BV785
CD3ε-AF700
CD4+ 85.3
CD4-BV711
CD3ε-AF700
CD44+ CD62L+ 74.1
CD44+ CD62L- 8.75
CD62L+ CD44- 13.9
CD44-PE-Cy7
CD62L-PE
CD44+ CD62L+ 83.5
CD44+ CD62L- 12.6
CD62L+ CD44- 2.65
CD44-PE-Cy7
CD62L-PE
c
FSC/SSC 81.1
SSC-A
FSC-A
Single Cells 93.3
FSC-H
FSC-A
Live CD45+ 66.3
CD45-BUV395
FVD700
Thy1+ 27.7
Thy1- 72.1
Thy1.2-FITC
CD11b-PacBlue
CD19+ 73.3
CD19- 26.3
CD19-PEDazzle594
CD11b-PacBlue
Eosinophils 0.21
SiglecF- 99.7
SiglecF-BV786
CD11b-PacBlue
Neutrophils 17.3
Ly6G- 82.3
Ly6G-APC-Cy7
CD11b-PacBlue
NK1.1- 91.5
NK1.1-BV650
CD11b-PacBlue
Monocytes 72.4
CCR2-BV711
CD11b-PacBlue
Ly6Chi 12.6
CD64+ 61.5
CD64- Ly6C- 24.4
Ly6C-BV605
CD64-AF647
Macrophages 94.5
MERTK-PCP-eF710
CD64-AF647
cDCs 23.1
MHCII-BUV737
CD11c-PE-Cy7
cDC2 20.1
cDC1 66.6
SIRPα-PE
CD11b-PacBlue
d
FSC/SSC 69.6
SSC-A
FSC-A
Single Cells 96.6
FSC-H
FSC-A
Live CD45+ 5.00
CD45-BUV395
FVD700
CD45-hi 23.8
Microglia 72.9
CD45-BUV395
CD11b-FITC
Neutrophils 34.2
Ly6G- 65.0
Ly6G-APC-Cy7
CD11b-FITC
61.9
SAM 27.9
8.61
1.64
TREM2-APC
CD9-BV421
cDC1 54.5
cDC2 45.5
MHCII-BUV737
CD11b-FITC
cDCs 2.34
MHCII-BUV737
CD11c-BV650
Monocytes 21.4
CCR2-BV711
CD11b-FITC
Ly6C-hi 42.8
CD64- Ly6C- 43.8
CD64+ 11.3
Ly6C-PerCP-Cy65.5
CD64-BV605

**Supplementary Fig. 1: Representative flow cytometry gating.**

**a-d,** Gating of T cells in cortical tissue (**a**), T cells in *ex vivo* coculture (**b**), myeloid cells in spleen (**c**), and myeloid cells in cortical tissue (**d**).
